# Reciprocal F1 hybrids of two inbred mouse strains reveal parent-of-origin and perinatal diet effects on behavior and expression

**DOI:** 10.1101/262642

**Authors:** Daniel Oreperk, Sarah A Schoenrock, Rachel McMullan, Robin Ervin, Joseph Farrington, Darla R Miller, Fernando Pardo-Manuel de Villena, William Valdar, Lisa M Tarantino

## Abstract

Parent-of-origin effects (POEs) in mammals typically arise from maternal effects or from imprinting. Mutations in imprinted genes have been associated with psychiatric disorders, as well as with changes in a handful of animal behaviors. Nonetheless, POEs on complex traits such as behavior remain largely uncharacterized. Furthermore, although perinatal environmental exposures, such as nutrient deficiency, are known to modify both behavior and epigenetic effects generally, the architecture of environment-by-POE is almost completely unexplored. To study POE and environment-by-POE, we employ a relatively neglected but maximally powerful POE-detection system: a reciprocal F1 hybrid population. We exposed female NOD/ShiLtJxC57Bl/6J and C57Bl/6JxNOD/ShiLtJ mice, *in utero*, to one of four different diets, then after weaning recorded their whole-brain gene expression, as well as a set of behaviors that model psychiatric disease. Microarray expression data revealed an imprinting-enriched set of over a dozen genes subject to POE; the POE on the most significantly affected gene, *Carmil1* (a.k.a. *Lrrc16a*), was validated using qPCR in the same and in a new set of mice. Several behaviors, especially locomotor behaviors, also showed POE. Interestingly, Bayesian mediation analysis suggests *Carmil1* expression suppresses behavioral POE, and *Airn* suppresses POE on *Carmil1* expression. A significant diet-by-POE was observed on one behavior, one imprinted gene, and over a dozen non-imprinted genes. Beyond our particular results, our study demonstrates a reciprocal F1 hybrid framework for studying POE and environment-by-POE on behavior.

## INTRODUCTION

It is well established that susceptibility to psychiatric disorders arises from a combination of genetics and environmental exposures (Lee and Avramopoulos 2014). Less well-studied is the phenomenon that this susceptibility seems to vary depending on whether certain harmful alleles were carried by the mother — or by the father (Davies *et al.* 2001; Isles and Wilkinson 2000). That is, it is unclear to what extent the heritable component of disease risk is driven by parent-of-origin effects (POEs). Especially poorly understood is the extent to which POEs depend upon environmental context during development, and therefore how alternate environmental exposures could modulate a POE’s impact on disease risk. A better understanding of POEs and their environmental modifiers could lead to improved interpretation of existing studies, to more effective experimental design, and even to novel public health interventions. Nonetheless, rigorous estimation of POEs in humans is difficult, especially on complex traits; even in animals it requires specialized experimental design attuned to POE biology.

Hager *et al.* (2008) used “parent-of-origin-dependent effect" to describe any genetic effect that causes phenotypic differences in reciprocal heterozygotes. Similarly, here we use “POE” as a shorthand for any effect driven by the interaction of genetic background with either: 1) maternal factors, encompassing maternal behaveior (Peripato and Cheverud 2002), oocyte composition (Tong *et al.* 2000), and *in utero* environment (Cowley *et al.* 1989; Kirkpatrick and Lande 1989); or 2) imprinting (Georges *et al.* 2003; Vrana *et al.* 2000; Wolf *et al.* 2014; Schultz *et al.* 2015), an epigenetic process in which either the maternally or paternally inherited allele of certain genes is at least partially silenced (Crowley *et al.* 2015; Bartolomei and Ferguson-Smith 2011).

Imprinting-driven POEs may be particularly relevant to psychiatric disease since evidence suggests imprinted genes affect behavior as well as brain development and function. In particular, many imprinted genes are active (some exclusively) in the brain (Prickett and Oakey 2012), especially during embryogenesis (Wilkinson *et al.* 2007); in mice, imprinted genes have been identified that affect brain size and organization (Wilkinson *et al.* 2007), control of nutritional resources (Bartolomei and Ferguson-Smith 2011), maternal nesting, pup-gathering/grooming, suckling, exploratory behavior, and social dominance (Dent and Isles 2014); and, in humans, imprinted gene mutations have been implicated in Prader-Willi and Angelman syndromes (Dykens *et al.* 2011), and in schizophrenia (Francks *et al.* 2007; Linhoff *et al.* 2009).

Imprinted genes may present an ideal path for understanding the interaction of genetics and environmental exposures — especially diet — on development: not only can imprinting be developmental-stage (and tissue)-specific (Koerner *et al.* 2009; Prickett and Oakey 2012), but it is also believed to largely result from differential allelic methylation, and thus to require dietary methyl donors (Crider *et al.* 2012). Moreover, rodent and human studies have demonstrated that certain perinatal diets affect both imprinting and behavior: for example, perinatal protein deficiency (PD) and vitamin D deficiency (VDD) both induce methylation changes (Vucetic *et al.* 2010; Lillycrop *et al.* 2007; Kesby *et al.* 2010, and alter behaviors that model schizophrenia (Burne *et al.* 2004a,b, 2006; Palmer *et al.* 2008; Franzek *et al.* 2008; Kesby *et al.* 2006, 2010; Burne *et al.* 2006; Harms *et al.* 2008, 2012; Turner *et al.* 2013; Vucetic *et al.* 2010). Similarly, other perinatal diets that imply a deficiency in methyl donors have been linked to reduced methylation in the brain (Davison *et al.* 2009; Niculescu *et al.* 2006; Konycheva *et al.* 2011), increased anxiety-like behaviors (Ferguson *et al.* 2005; Konycheva *et al.* 2011), and changes in learning and memory (Konycheva *et al.* 2011; Berrocal-Zaragoza *et al.* 2014). In general, epigenetic effects have repeatedly been shown to be sensitive to maternal diet during the prenatal period: classically in Agouti mouse experiments (Waterland and Jirtle 2003); and observationally in studies of human physiology, mental health, and gene expression during the Dutch Hunger Winter (Heijmans *et al.* 2008; Tobi *et al.* 2009).

### Reciprocal F1 hybrids (RF1s) for investigating POE and its environmental modifiers

The above findings motivate an experiment to directly determine the extent of POEs on psychiatric disease across multiple perinatal dietary exposures in a simple, controlled, and replicable system — something only possible in an animal model. An ideal population is provided by (female) reciprocal F1 hybrids (RF1s) of inbred strains: in female RF1s, genetic background is constant (save for mitochondria), and only the direction of inheritance varies, allowing POEs to be measured directly. RF1s have been used to identify POEs on behavior in a handful of studies so far (Putterman 1998; Isles *et al.* 2001; Calatayud and Belzung 2001; Calatayud *et al.* 2004). Here we exploit the replicability of RF1s further to study the unconfounded effect of an environmental modifier on POE, varying diet while genetic background stays constant. To our knowledge, this approach has only been followed in Schoenrock *et al.* (2017), our recent related study in which we reciprocally crossed Collaborative Cross strains.

Here we examine, under four different *in utero* dietary exposures, behavior and expression in RF1s of the inbred mouse strains C57BL/6J (B6) and NOD/ShiLtJ (NOD). Mouse was chosen because of its rapid gestation and development, its versatility as a model for behavioral genetics and environmental perturbation, and its similarity to humans in functional consistency of imprinting (Bartolomei and Ferguson-Smith 2011). B6 and NOD inbred strains were selected because: 1) B6 is the reference genome and is the best characterized strain with respect to behavior; 2) B6 and NOD are among the founder strains for the Collaborative Cross, a population that is an area of focus for our labs; 3) B6 and NOD were both readily available, and B6-NOD crosses generate large litters, facilitating replication; 4) NOD is genetically similar enough to B6, that standard B6-expression microarrays were appropriate for NOD alleles as well (Oreper et *al.* 2017), but different enough that a substantial number of POEs on gene expression could still be revealed by B6-NOD RF1s.

Our replication of the RF1s under four different in *utero* dietary exposures serves several purposes, namely to: 1) increase the likelihood of observing POE, as POE may be diet-specific; 2) estimate the extent to which POE generalizes across alternate perinatal dietary exposures; and 3) estimate the perinatal diet effect itself.

Our study, the first to examine the connection between POE on expression and POE controlled behavior, demonstrates: 1) the presence of POEs on behavior and gene expression, many of which are robust to differences in perinatal diet; 2) a possible explanatory pathway connecting imprinting, to gene expression, to behavior; and 3) the usefulness of our approach as a template for further animal model studies of POE and developmental exposures on complex traits.

## EXPERIMENTAL MATERIALS AND METHODS

### Mice

C57BL/6J (B6) and NOD/ShiLtJ (NOD) mice originated from a colony maintained by Gary Churchill at Jackson Laboratory, and were transferred in 2008 to the FPMdV lab at UNC (this originating colony also produced the G1 breeders of the CC; see Srivastava et *al.* 2017). Six-week old B6 females (3-8 dams/diet) and NOD females (3-5 dams/diet) were transferred from the FPMdV lab to the Tarantino lab at UNC and acclimated for one week. At 7 weeks of age, dams were placed on one of 5 different diets. At 12 weeks, dams were mated with males of the opposite strain to produce either B6xNOD or NODxB6 F1 hybrids (dam strain listed first; Figure 1B). Pregnant dams remained on their experimental diet until litters were weaned, ensuring that offspring were exposed to the diet throughout the entire perinatal period. At postnatal day (PND) 21, F1 hybrids were weaned onto a standard laboratory chow (Pico rodent chow 20; Purina, St. Louis, MO, USA) (Figure 1A). F1 hybrids were bred in one vivarium, but then transferred to a separate behavioral testing vivarium, and were then acclimated to this testing vivarium for at least one week before testing began. Mice were housed in a specific pathogen free facility on a 12-hour light/dark cycle with lights on at 7 A.M. All procedures and animal care were approved by the UNC Institutional Animal Care and Use Committee and followed the guidelines set forth by the National Institutes of Health (NIH) Guide for the Care and Use of Laboratory Animals.

**Figure 1.**
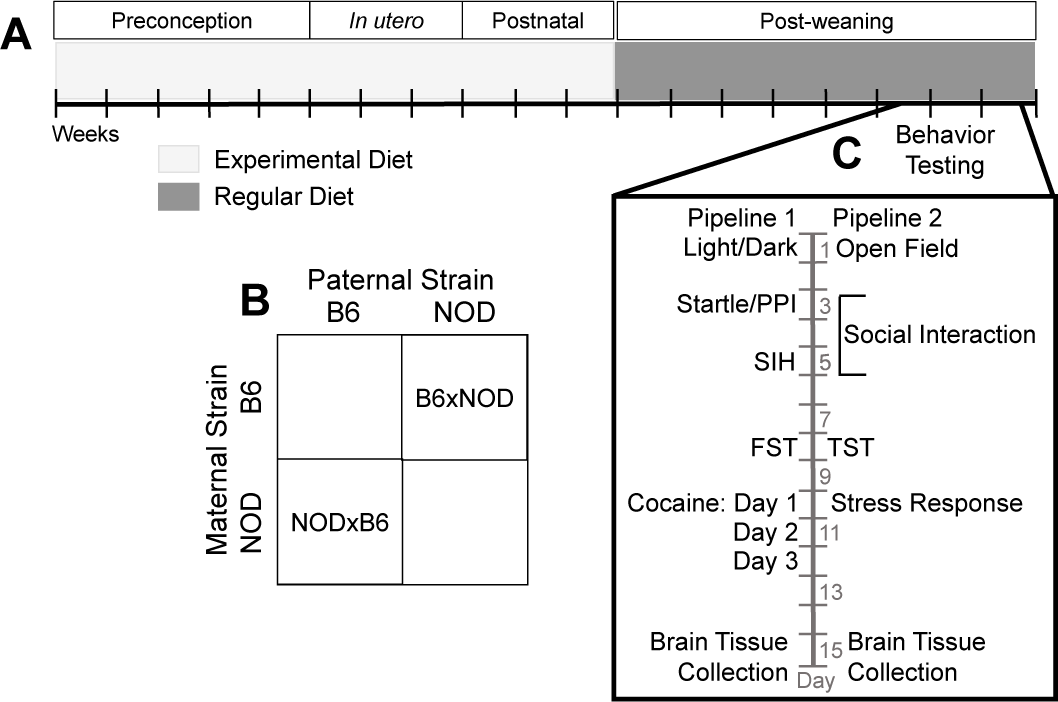
Experimental design to assess POE, perinatal diet, and diet-by-POE on behavior and gene expression in reciprocal F1 hybrids (RF1s). Female NOD/ShiLtJ (NOD) and C57BL/6J (B6) mice were placed on one of 4 experimental diets (protein deficient, vitamin D deficient, methyl enriched, standard) at 7 weeks of age (A). After 5 weeks, NOD females were mated to B6 males and B6 females to NOD males forming NODxB6 and B6xNOD RF1 hybrids, respectively (B). Dams remained on their experimental diet throughout gestation and the postnatal period. At PND 21, female F1 hybrids were weaned and placed onto a regular laboratory diet (A). Upon reaching adulthood at PND 60, F1 hybrids were tested in one of two behavioral pipelines. After behavioral testing, mice were euthanized, and their brain tissue collected for gene expression analysis via microarray and qPCR (C).

### Experimental Diets

The following diets, purchased from Dyets Inc. (Bethlehem, PA), were administered: vitamin D deficient (VDD; #119266), protein deficient (PD; 7.5% casein; #102787), methyl donor deficient (MDD; #518892), methyl donor enriched (ME; #518893) and control (Std; #AIN-93G). The PD and VDD diets were nutritionally matched to the Std diet while the MDD was matched to the ME diet; Table S1 specifies each diet’s nutrient composition. Food and water were available *ad libitum* throughout the experiment.

### Behavior Assays

To ensure a standardized genetic background that included the sex chromosomes, the only tested F1 hybrids were female. Mice were 61.7 days old ± 2.6 standard deviations at the onset of testing. All behavioral testing was performed during the light part of the light/dark cycle between 8:00 A.M. and 12:00 P.M. Mice were placed into one of two behavioral pipelines (Figure 1C) to assess anxiety- and depressive-like behavior, stress response, sensorimotor gating or response to a psychostimulant: Pipeline 1— light/dark assay, startle/prepulse inhibition (PPI), stress-induced hyperthermia (SIH), forced swim test (FST) and cocaine response (N = 91); Pipeline 2— open field (OF), social interaction test, tail suspension and restraint stress (N = 87). In total, 34 behavioral measures were collected, with 22 in pipeline 1 and 12 in pipeline 2 (Table 1). For each pipeline, mice were tested in 3 batches, over months. Offspring from both RF1 directions, as well as from at least 2 diet exposures, were included in each batch, to avoid confounding. For each diet and RF1 direction, we tested litters from at least 2 dams (N = 4 ± 1.4; see Table S2 for dam and offspring counts). One mouse in the NODxB6 ME group was euthanized due to injury on the day of social interaction testing; there is no data for this mouse for social interaction or for any subsequent test. There is no restraint stress data for another 4 mice (1 NODxB6 ME, 2 B6xNOD Std, 1 B6xNOD VDD), due to either death in the restrainer or insufficient serum collected for radioimmuno assay (RIA) analysis of corticosterone (CORT) levels.

**Table 1.**
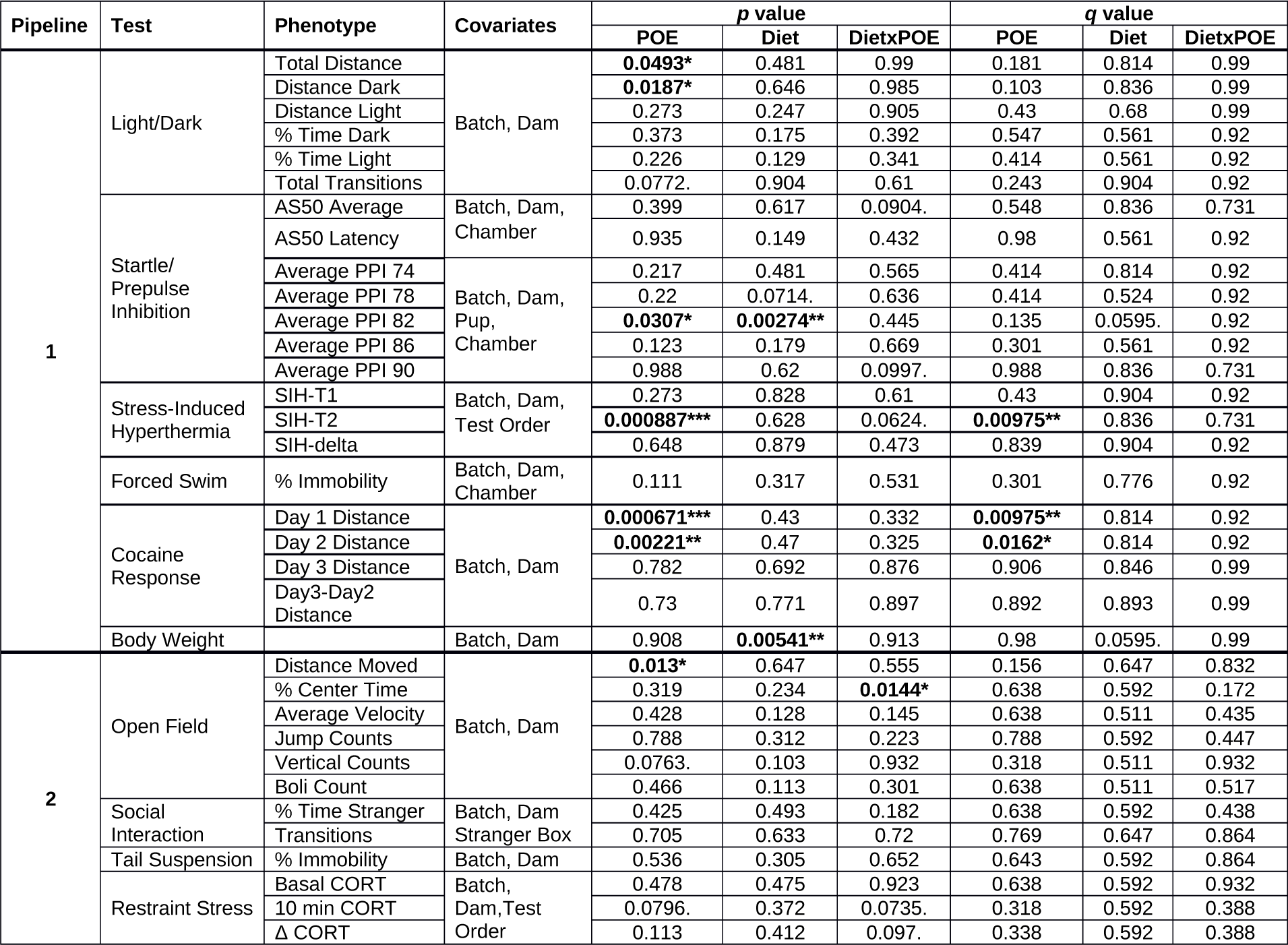
POE, perinatal diet effect, and diet-by-POE on behavioral phenotypes. For each phenotype, the table shows the modeled variables, along with the p-values of interest, and their corresponding q-values (FDR), which account for multiple testing within a behavioral pipeline. Significant values are **bolded**, and *, **, and ***, indicate significance levels of *0.05, **0.01, ***0.001 respectively. POE = parent of origin effect; PPI = prepulse inhibition; CORT = corticosterone; SIH-T1 = basal temperature; SIH-T2 = post-stress temperature; SIH-delta = (T2-T1)

**Table 2** Microarray-measured effects on expression. For each effect type/significance threshold type, the table specifies the significance threshold value, as well as the number of probesets, genes, and imprinted genes whose expression was significantly affected. Note that: i) some probesets measure multiple genes, and some genes are measured by multiple probesets; ii) the FDR and FWER thresholds for diet differ greatly; iii) imprinting is enriched among genes subject to POE, and iv) by FWER, diet does not affect any imprinted gene, whereas one imprinted gene is subject to diet-by-POE

#### Open Field (OF)

Mice were placed in the OF arena for 10 minutes. The OF apparatus (ENV-515-16, Med Associates, St. Albans, VT, USA) was a 43.2x43.2x33 cm arena, consisting of a white Plexiglas floor and clear Plexiglas walls with infrared detection beams at 2.54 cm. intervals on the x, y, and z axes that automatically tracked mouse position and activity throughout each experimental session. The apparatus was in a sound-attenuating chamber (73.5x59x59 cm) fitted with two overhead light fixtures containing 28-V lamps. Mice were placed in the OF arena for 10 minutes. The OF apparatus (ENV-515-16, Med Associates, St. Albans, VT, USA) was a 43.2x43.2x33 cm arena, consisting of a white Plexiglas floor and clear Plexiglas walls with infrared detection beams at 2.54 cm. intervals on the x, y, and z axes that automatically tracked mouse position and activity throughout each experimental session. The apparatus was in a sound-attenuating chamber (73.5x59x59 cm) fitted with two overhead light fixtures containing 28-V lamps. Mice were scored for total distance traveled (cm), average velocity (cm/s), number of vertical movements (rearing), and percent time spent in the center of the arena (a 22.86 cm^2^ central part of the arena). These measurements were recorded in 5 bins of 2-minute width, and were scored in post-session analyses using Activity Monitor 5.1 software (Med Associates). The testing apparatus was cleaned with a 0.25 % bleach solution between test subjects.

#### Social Interaction

Social approach was measured in a 3-chamber social interaction apparatus during a 20-minute test (described fully in Moy *et al.* (2007)). Briefly, the first 10 minutes was a habituation period in which the test mouse was given free access to all 3 chambers. The total number of transitions between all chambers during this 10 min period was measured. During the second 10 minutes, the test mouse was given the choice between a chamber containing a circular mesh enclosure that held a stranger mouse (B6), and a chamber containing an empty mesh enclosure. The amount of time the test mouse spent in the chamber with the stranger mouse was recorded and is reported as “percent stranger time”, a measure of social preference.

#### Tail Suspension

Mice were suspended by a piece of laboratory tape wrapped around the tail and hung from a hook at the top of a 24.13 cm × 17.78 cm × 17.78 cm white acrylic enclosure. Mice were videotaped for the entire 4-minute session, and videotapes were analyzed for immobility during the last 2 minutes using the Actimetrics Freeze Frame analysis program (Actimetrics, Wilmette, IL). Percent immobility during the last two minutes is reported as a measure of depressive-like behavior (Miller *et al*. 2010).

#### Restraint Stress

Restraint was used to elicit a stress response that was then quantified by measurement of CORT levels in the serum. A retroorbital blood sample was taken immediately prior to placing the mice into a Broome-Style restraint tube (Plas Labs, Inc., Lansing, MI, USA) for 10 minutes. Immediately upon removal from the restrainer, a second retroorbital eye bleed was performed. Whole blood was centrifuged to isolate serum, and then the CORT levels were measured by competitive RIA per the manufacturer’s protocol (MP Biomedicals, Santa Ana, CA, USA).

#### Light/Dark

The open field arena described above was converted to a light dark apparatus by placement of an opaque polycarbonate black box that occupied one third of the arena space, thus allowing the mouse to choose between the light or dark part of the apparatus. Mice were placed in the lighted area immediately adjacent to and facing the entry to the dark enclosure and remained in the apparatus for 10 minutes. The amount of time (sec), distance moved (cm) and number of transitions between the dark and light zones was scored in 5-minute bins in post-session analyses using Activity Monitor 5.1 software (Med Associates). The testing apparatus was cleaned with a 0.25 % bleach solution between test subjects.

#### Startle and prepulse inhibition (PPI)

Acoustic startle and PPI of the startle response were both measured using the San Diego Instruments SR-Lab system (San Diego, CA), and following the protocol in Moy *et al*. (2012). Mice were placed in a plexiglas cylinder located in a sound-attenuating chamber that included a ceiling light, fan, and a loudspeaker that produced the acoustic stimuli (bursts of white noise). Background sound levels (70 dB) and calibration of the acoustic stimuli were confirmed with a digital sound level meter. Each test session consisted of 42 trials, presented following a 5-min habituation period. There were 7 types of trials: no-stimulus trials, trials with a 120 dB acoustic startle stimulus (a.k.a., ASR), and 5 trials in which a 20 ms prepulse stimulus (74, 78, 82, 86, or 90 dB) was presented 100 ms before the onset of the 120 dB startle stimulus. The different trial types were presented in 6 blocks of 7, in randomized order within each block, with an average intertrial interval of 15 sec (range: 10 to 20 s). Measures were taken of the startle response amplitude (RA) for each trial, defined as the peak response recorded from the onset of startle stimulus to the end of the 65-msec sampling. The PPI for each prepulse sound level was calculated as:

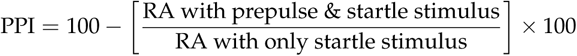

#### Stress-induced hyperthermia (SIH)

Each tested mouse was individually removed from its home cage, and then its body temperature (T1) was measured. Specifically, a lubricated digital thermometer probe was inserted 1-1.5 cm into the rectum for approximately 10 seconds. The mouse was then returned to its home cage, and 10 minutes later the temperature measurement was repeated (T2). The difference in body temperature, ΔT = T2 – T1, was used as a measure of anxiety-like behavior. Basal temperature was measured for all mice within a single cage in under a minute, to avoid increases in body temperature due to anticipatory stress.

#### Forced swim test (FST)

Mice were placed in a glass-polycarbonate cylinder (46cm tall ×21cm in diameter) filled with water (25-28 °C) to a depth of 15 cm for 6 minutes. The duration of immobility during the last 4 minutes of the test period was scored using Ethovision 7.0 automated tracking software (Noldus, Leesburg, VA). Immobility was defined as no movements other than those needed to stay afloat. Mice were monitored continuously, and removed if they were unable to keep their nose or heads above water for more than 30 seconds. Percent immobility was reported as a measure of depressive-like behavior.

#### Cocaine-induced locomotor activation

Cocaine-induced locomotor activity was measured over a 3-day test protocol in the OF arena described above. On days 1 and 2, mice were given an intraperitoneal injection of saline before being placed into the OF arena for 30 minutes, and then returned to their home cage. The day 3 protocol was nearly identical, but instead of saline, mice were injected with 20 mg/kg cocaine (Cocaine HCl; Sigma-Aldrich, St. Louis, MO). The total distance traveled was calculated for each day, and cocaine-induced locomotor activation was calculated by subtracting the distance on day 2 from day 3.

#### Body Weight

Adult body weight was recorded for mice in pipeline 1 prior to startle/PPI and cocaine administration.

### Gene Expression

To identify genes subject to POE and/or perinatal-diet effect, whole-brain expression was measured by microarray, and key expression results were later validated with qPCR.

#### Tissue extraction

Three days after completion of behavioral testing, mice were euthanized, cerebellar tissue was removed, and the brain was split midsagitally into left and right hemispheres. Brain tissue was flash frozen in liquid nitrogen. Right brain hemispheres were pulverized using a BioPulverizer unit (BioSpec Products, Bartlesville, OK). Pulverization batches were designed to prevent contamination between mice from different crosses or diets.

#### RNA extraction

Total RNA was extracted from 25 mg of powdered brain hemisphere tissue using an automated bead-based capture technology (Maxwell 16 Tissue LEV Total RNA Purification Kit, AS1220; Promega, Madison, WI). Purified mRNA was evaluated for quality and quantity by Nanodrop Spectrophotometer (Thermo Scientific).

#### Microarray expression measurement

Of the 178 behaviorally-phenotyped, female B6xNOD and NODxB6 F1s, 96 females were selected for microarray measurement of gene expression. The choice of 96 mice was balanced to include both directions of reciprocal cross offspring, all 4 diets, as well as both behavioral test pipelines, while simultaneously maximizing the number of represented litters. Gene expression was measured using the Affymetrix Mouse Gene 1.1 ST Array. All samples were processed by the Functional Genomics Core at UNC.

#### qPCR expression measurement

Commercially available Taqman qPCR assays for *Carmil1* (Life Technologies, Mm01158156_m1) and *Meg3* (Life Technologies, Mm00522599_m1) were used to estimate gene expression levels. For each sample, mRNA was retro-transcribed to cDNA using 200ng of starting RNA (SuperScript III First-Strand Synthesis System, 18080051; ThermoFisher Scientific, Waltham, MA) following the manufacturer’s protocol. The amplification curve was calibrated using an *Rfng* (Life Technologies, Mm00485703_m1) reference assay. All assays were performed following the manufacturer’s protocol on an ABI StepOne Plus Real-Time PCR System (Life Technologies, Carlsbad, CA), and in duplicate; each sample was assayed on 2 of 3 available plates. Samples were plated such that breeding batch, which explained much of the microarray expression variance, was partially confounded with qPCR plate. Cycle thresholds were determined using ABI CopyCaller v2.0 software on default settings. All available brain samples were assayed, regardless of hemisphere.

## COMPUTATION AND STATISTICAL MODELS

### Statistical Analysis of Behavior

Diet effects, POE, and diet-by-POE were evaluated using a mostly similar linear mixed model (LMM) for every behavior. Specifically, each behavioral phenotype was transformed to ensure residual normality (Appendix B), and then modeled by an LMM that: 1) controlled for batch and any test-specific nuisance factors; 2) controlled for population structure by modeling dam as a random effect; and 3) modeled diet, PO, and diet-by-PO using categorical fixed effects. See Appendix A for more details.

Every behavioral LMM was fit in R (R Core Team 2016) using lme4 (Bates *et al.* 2015) and *p*-values calculated by a type I (i.e., sequential) sum of squares ANOVA using Satterthwaite’s approximation using lmerTest (Kuznetsova *et al.* 2015). To account for multiple testing, the *p*-values were pooled over all behaviors in each pipeline, but separated per effect type (diet effects, POE, diet-by-POE); then, each pipeline/effect type combination was subject to a Benjamini-Hochberg false discovery rate correction, generating q-values (Benjamini and Hochberg 1995).

Most test-specific nuisance factors were modeled as fixed effects, including: 1) the swimming chamber for the forced swim test; 2) the testing order for the stress induced hyperthermia and restraint stress tests; and 3) the box holding the stranger mouse for the sociability test. In repeated measures models of the startle/PPI phenotypes, random effects were used for pup and chamber (Appendix A).

For ASR data, the modeled outcome was the raw ASR divided by the mouse body weight. For the PPI at each prepulse intensity, the modeled outcome was the average PPI response divided by the weight-adjusted ASR value— a weight-and-ASR-adjusted PPI.

### Microarray Preprocessing

Microarray probe alignments to the GRCm38.75 C57BL/B6J reference genome (the reference we use throughout) were used to infer probe binding locations (Appendix G). Using these locations, along with Affymetrix Power Tools (APT) 1.18 software (Affymetrix 2017), probes and probesets at biased/uninformative binding locations (Appendix H) were masked. Masking reduced the original set of 28,440 non-control probesets to only 20,099 probesets (representing 19,224 unique genes, including X chromosome genes). For each remaining probeset, RMA (Irizarry *et al.* 2003) was applied to the non-masked probes to compute a probeset expression score. Each probeset’s position was defined as the binding location of its first non-masked probe. The expression of one mouse was inadvertently measured twice; these probeset measurements were pairwise averaged.

### Statistical Analysis of Gene Expression

Data from 95 microarray-assayed mice and 20,099 probesets was used to test diet effects, POE, and diet-by-POE on gene expression as follows. For each probeset: 1) fixed nuisance effects were regressed out of the expression score to generate adjusted expression values (see below); 2) the adjusted expression was transformed to ensure residual normality; 3) the resulting values were tested for diet, POE and diet-by-POE using an LMM that accounted for dam (using the R package nlme Pinheiro *et al.* 2016).

The p-value distribution for each effect type appeared to be inflated. To correct for the inflation, p-values were adjusted by a genomic-control-like procedure (Dadd *et al.* 2009) whereby, for all p-values within an effect-type, an inflation factor was estimated and then divided out (Appendix E). Then, to control for multiple testing, we used two complementary approaches: Benjamini-Hochberg false discovery rate (FDR; Benjamini and Hochberg 1995), applied separately per effect type; and family-wise error rate (FWER) control, using a bespoke permutation procedure that makes minimal parametric assumptions while accounting for between-probeset correlations (Appendix F).

#### Adjusted expression, SSVA estimated nuisance factors

Prior to testing for diet effects, POE and diet-by-POE, expression values for each probeset were first adjusted by regressing out nuisance effects; this was done to facilitate permutation-based threshold calculation (Appendix F). Nuisance effects were estimated by fitting a simple linear model (to the original expression) that accounted for nuisance factors only — batch, pipeline, and a set of estimated unobserved factors. These unobserved factors were themselves estimated using a modified form of Supervised Surrogate Variable Analysis (Leek 2014), which we adapted to accommodate random effects (Appendix D).

### Analysis of imprinting status

Each microarray probeset was classified as measuring imprinted gene expression, if its probe sequences either: 1) hybridized to the sequence of an imprinted gene identified in Mousebook (Blake *et al*. 2010) or in Crowley *et al*. (2015); or 2) hybridized within 100bp of these known imprinted genes. All together, 241 probesets were classified as measuring imprinted regions, corresponding to 182 unique imprinted genes. Each probeset was also categorized as to whether it revealed a significant (q-value < 0.05) POE on expression of the probed region. Fisher’s exact test was used to calculate p-values for the association between imprinting status and significant expression POE.

### Analysis of qPCR validation data

An apparent POE on microarray expression of *Carmil1* and a diet-by-POE on *Meg3* were validated by analysis of their respective qPCR data as follows. Each gene’s qPCR relative-cycle-threshhold (relative to *Rfng*, Appendix I) was transformed for residual normality, and then modeled by an LMM that accounted for pipeline, the interaction of breeding batch with qPCR plate (as a random efffect), and dam (random effect), as well as the diet, POE, and diet-by-POE effects. LMMs were fit using lme4 (Bates et *al.* 2015), with p-values computed using lmerTest (Kuznetsova et *al.* 2015). qPCR data analysis was repeated in three sets of mice: 1) 85 mice assayed by both microarray and qPCR; 2) 30 mice newly assayed by qPCR alone; and 3) all 115 qPCR’d mice.

### Mediation Analysis

POEs were observed upon several behaviors, as well as upon the expression of the non-imprinted gene, *Carmil1*. To identify (potentially imprinted) genes exerting POE on these outcomes, we applied a genomewide mediation analysis. That is, for each outcome above, and for each potential mediator gene, we tested whether the gene’s expression mediated POE on the outcome (details in Appendix J). For completeness, and to generate percentile-based significance thresholds, we tested every gene as a candidate mediator whether or not we observed POE on the candidate in mediation-free analysis.

This test was performed using a model (see Figure 8 notation) in which the outcome was the sum of: 1) outcome-specific nuisance effects (which also affect the candidate mediator gene); 2) a diet-specific *direct* effect of parent-of-origin 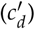, and 3) a diet-specific *indirect* effect of parent-of-origin, that is mediated by way of POE on the candidate mediator gene’s expression (*a*_*d*_*b*). Candidate mediator genes with a significant *average* indirect effect 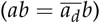 on POE were identified as true mediators. Candidate mediator genes for which the indirect and direct effect had opposite signs were further classified as suppressors.

We note that in this model, diet does *not* modulate the effect of mediator expression on outcome; the indirect effect is diet-specific only insofar as diet affects mediator expression.

#### Mediation analyzed using a Bayesian approach

Most simple mediation analyses are handled using frequentist methods. However, our mediation model required that we estimate an indirect effect across multiple diets, all while accounting for the random effect for dam. For this type of complexity, a Markov Chain Monte Carlo (MCMC)-based Bayesian approach was ideal, providing the necessary flexibility to easily provide point and interval estimates of the indirect effect, all without the need to derive an analytic form (Yuan and MacKinnon 2009; Wang and Preacher 2015). Our mediation model, described in more detail in Appendix J, was implemented in JAGS [Just Another Gibbs Sampler; Plummer (2003, 2016)]. Posterior medians and credible intervals for direct and indirect effects were estimated from Gibbs samples. To obtain a measure of “mediation significance”, we estimated the indirect effect’s “Combined Tail Probability” (CTP): the minimum of the sample-based, upper and lower tail probabilities of the indirect effect, where we deemed CTP ≤ .05 significant (as used in, *e.g.*, Schoenrock *et al.* 2016).

#### Mediation of *Carmil1* expression

Mediation modeling of the *Carmil1* expression outcome was restricted to data from mice in which expression was measured. Batch, pipeline, and dam (a random effect), were modeled as nuisance effects acting on both the mediator gene and on *Carmil1.*

#### Mediation of behavior

All behavior outcomes were tested for gene mediation of POE, whether or not expression-free analysis had revealed POE on that outcome. Modeling was restricted to data from mice in which expression and behavior were both measured. Dam, batch, and behavior-specific covariates were modeled as nuisance effects on both mediator and outcome. Pipeline was *not* modeled, as each behavior was only measured in one pipeline. For PPI outcomes, groups of measurements from the same mouse/prepulse intensity were averaged together into a single value.

#### Aggregate mediation of behavior

To quantify each gene’s aggregate level of mediation over *all* behaviors, we defined a statistic inspired by the Fisher combined p-value (Fisher 1925): the “Combined Tail Probability" (CTP; Appendix J). Aggregate levels of mediation were also assessed by counting how often a given mediator was among the 3 most significant mediators for any behavior.

### Reporting significant genes vs. probesets

The number of genes we report as significantly affected by some factor (*e.g.*, diet) is generally not equal to the number of significantly affected probeset measurements. The mismatch arises because some genes (*e.g., Snord 115)* are assayed by more than one probeset, and some probesets simultaneously assay more than one gene (*e.g.*, overlapping genes). For each significantly affected multi-gene probeset, we propagate significance to all of its assayed genes.

### Test for miRNA regulation of significantly affected genes

To evaluate the validity of the diet-by-POE on *Mir341*, we tested whether the set of other genes showing diet-by-POE (by FDR) was enriched for *Mir341’s* predicted targets of regulation. Specifically, we used miRHub (Baran-Gale *et al.* 2013), allowing it to consider all miRNA targets predicted by TargetScan (Agarwal *et al.* 2015), regardless of whether those targets were conserved in another species.

### Segregating variant determination

Variants segregating between NOD and B6 with > .95 probability were identified using ISVdb (Oreper *et al.* 2017).

### Computational resources

Computation was performed on Longleaf, a slurm based cluster at UNC. Up to 400 jobs were run at a time in parallel. Computation completed in approximately 6 days.

### Data Availability

Data and supplemental results files are stored on Zenodo at https://doi.org/10.5281/zenodo.1168578 (Oreper *et al.* 2018). File S1 contains detailed descriptions of all supplemental files. File S2 contains chromosome sizes. File S3 contains exon data. File S4 contains Snord data. Files S5, S6, and S7, contain imprinted genes from Crowley *et al.* (2015), Mousebook, and the union thereof, respect-tively. File S8 contains NOD variants. File S9 contains covariates for RF1s. Files S10 and S11 contain Affymetrix library files for the Exon 1.1 ST and 1.0 ST microarrays, respectively. File S12 contains 1.0 ST probe binding locations. File S13 contains raw (CEL) microarray-measured expression for RF1s. File S14 contains a summary of microarray expression— the output from APT-summarize, but with default args. File S15 contains pulverized brain data, pre-qPCR validation. File S16 contains qPCR data. File S17 contains behavior models for mediation analysis. File S18 contains body-weights. File S19 contains cocaine responses. File S20 contains FST data. File S21 contains light/dark data. File S22 contains OF data. File S23 contains restraint stress data. File S24 contains SIH data. File S25 contains sociability data. File S26 contains startle/PPI data. File S27 contains tail suspension data. File S28, S29, and S30 contain POE, diet, and diet-by-POE expression modeling results, respecttively. File S31 and S32 contain mediation analysis results for the *Carmil1* and behavior outcomes, respectively. Code to generate results is available at https://github.com/danoreper/mnp2018.git.

## RESULTS

### Overview and key results

NOD and B6 mice were reciprocally crossed, with F1 hybrids exposed perinatally to Std, VDD, ME, MD, and PD diets (the MD diet was eventually dropped due to a near total lack of reproductive/weaning productivity; Table S2). Following weaning, the female F1 hybrids were tested in one of two different pipelines, each of which consisted of a different set of behavioral tests (Figure S1). Following behavioral testing, whole brain gene expression was measured via microarray. Analysis and validation lead to the following key results (detailed in subsequent subsections):

- Parent-of-origin affected 7 behaviors, including multiple locomotor behaviors and SIH behavior.
- Perinatal diet affected body weight and PPI behavior.
- Diet-by-POE acted on OF percent center time.
- Diet, POE, and diet-by-POE significantly (by FWER) acted on expression of 37, 15, and 16 genes respectively.
- The significance of diet’s effect on expression was primarily driven by ME.
- Notable POE were observed on *Snord 115, Airn*, and most significantly on *Carmil1*, a non-imprinted gene.
- The *Carmil1* POE was qPCR-validated in two sets of mice: the microarrayed mice, and a new set of mice.
- Genes affected by POE are enriched for imprinting.
- POE on *Carmil1* seems to be mediated (specifically, suppressed) by the expression of the imprinted gene *Airn*;
- *Carmil1*, and *Snord 115*, and especially *Airn* seem to mediate POE on multiple behaviors. These, along with other identified mediators of behavioral POE, tend to be suppressors.

### Effects on behavior

At a nominal level, POE, diet, and diet-by-POE acted significantly upon 7,4, and 2 behaviors, respectively. Post-FDR correction, POE, diet, and diet-by-POE acted upon 3,0, and 0 behaviors, respectively. Table 1 shows per-variable p-values, whereas Table S5 shows tukey p-values for variable level contrasts.

#### POE acts upon several locomotor behaviors, as well as SIH and PPI outcomes

Across several assays and both pipelines, a significant POE was observed on 5 different assessments of locomotor behavior. In all 5 assesments, NODxB6 mice moved more than B6×NOD mice. In pipeline 1, in the Light/Dark test, a POE was observed on both total distance and distance moved on the dark side of the arena (p=0.0493, q=0.181; p=0.0187, q=0.103 respectively), but not on light side distance (p=0.273; Figure 2A). Also in pipeline 1, in the cocaine response assay, a POE was observed on total OF distance, on both the baseline and the habituation day (Day 1, p=0.000671, q=.00975; Day 2, p=0.00221, q=0.0162 respectively) (Figure 2B). In pipeline 2, in a separate set of OF-assessed mice, a POE was observed upon total-distance moved (p=0.013, q=0.156; (Figure 2B).

**Figure 2.**
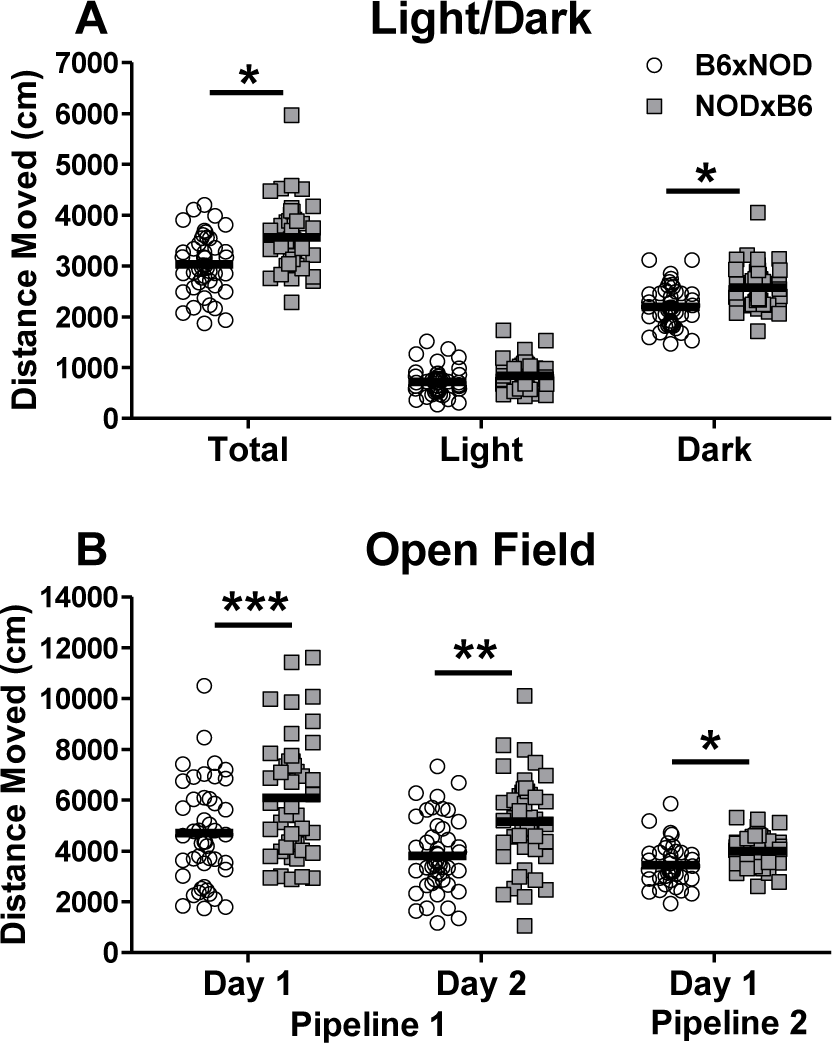
POEs on locomotor behavior are consistent across behavioral tests and pipelines. (A) Light side, dark side, and total distance moved in the light/dark arena for individual B6×NOD (n=46) and NOD×B6 (n=45) mice (bars indicate mean). (B) OF distance moved for B6×NOD (n=46) and NOD×B6 (n=45) mice, in Pipeline 1 on Day 1 and 2 of a 30 min cocaine response test when the mice received an ip saline injection; Distance moved in Pipeline 2 in a separate 10 min OF test (B6×NOD:n=39, NOD×B6:n=48). For all assays, NOD×B6 mice move significantly more than B6xNOD mice. *p <0.05, **p< 0.01, ***p< 0.001

POE was also observed on post-stress temperature in the SIH assay, with B6×NOD mice having higher temperatures (p=0.000887, q=0.00975; Figure 3). A smaller, non-significant effect in the same direction was also seen for both basal temperature (SIH-T1) and change in temperature (SIH-delta), consistent with a small difference in basal temperature being magnified after stress. A signify-cant POE was also observed on PPI at 82 decibels, with B6×NOD mice exhibiting a higher percent PPI than NOD×B6 (p=0.0307 and q=0.00274; Figure S2A). A similar effect (to that at 82 decibels) was observed at 86 decibels, but it was not significant (Figure S2A).

**Figure 3.**
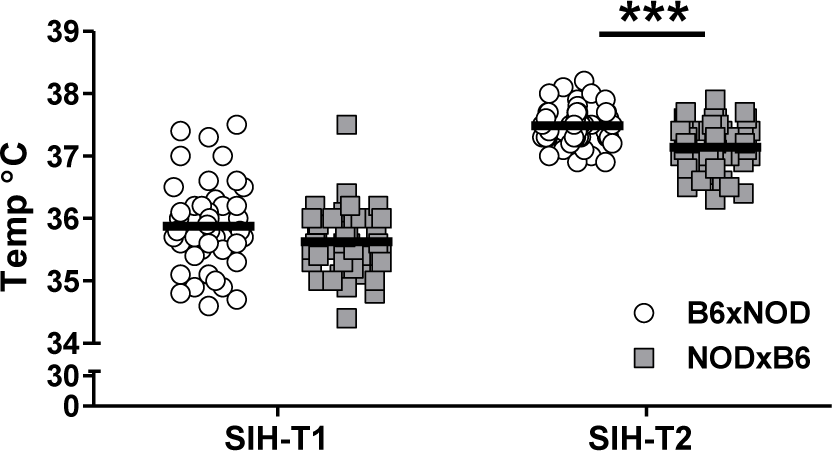
POEs on baseline (SIH-T1) and post-stress induced temperature (SIH-T2) in the stress induced hyperthermia test. Data are for individual B6xNOD (n=46) and NOD×B6 (n=45) mice (bars indicate mean). For SIH-T2 B6×NOD mice have higher temperature than NOD×B6 mice. A similar, though non-significant pattern seems to occur in the the SIH-T1 data

#### Diet has nominally significant effects on body weight and PPI

At a nominal level, perinatal diet significantly affected body weight (p=0.00541, q=.0595; Figure 4), with mice exposed to ME diet weighing less than mice exposed to Std and VDD diets (Tukey post-hoc p=0.0228 and p=0.0402). Diet also significantly affected measures of sensorimotor gating: in particular, PPI at 82 decibels (p=0.00274, q=0.0595; Figure S2B). At 78 decibels, PD had a non-significant (p=.0714, q =.524), but similar effect (Figure S2B). At both 78 and 82 decibels, PPI seemed greatest for PD mice compared to other diets, although individual contrasts were not significant (Figure S2B, Table S5)

**Figure 4.**
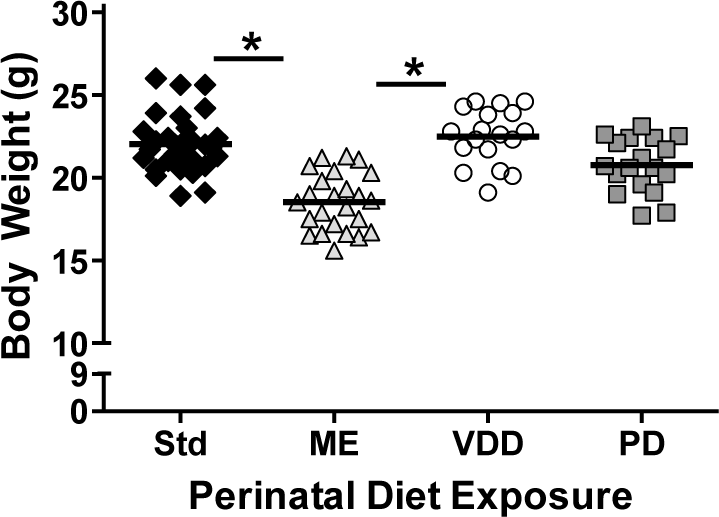
Effect of perinatal diet exposure on body weight in adulthood. Body weight of individual mice (bars indicate mean) exposed to either standard (Std, n=31), methyl enriched (ME, n=24), protein deficient (PD, n=18) or vitamin D deficient (VDD, n=18) diet duriug the perinatal period. Perinatal diet significantly affected body weight (p=0.00541). *indicates a significant difference between ME from Std and VDD mice (p<0.05)

#### Diet interacts with parent-of-origin to alter percent center time

A nominally significant diet-by-POE was observed on percent center time in the OF test (p=0.0144, q=0.172; Figure 5). In this test, NOD×B6 mice exposed to VDD and PD diets spent more time in the center of the arena than diet-matching B6×NOD mice, but no such difference was seen for ME or Std diets. Similar but non-significant effects were seen on OF locomotor activity (Figure S3).

**Figure 5.**
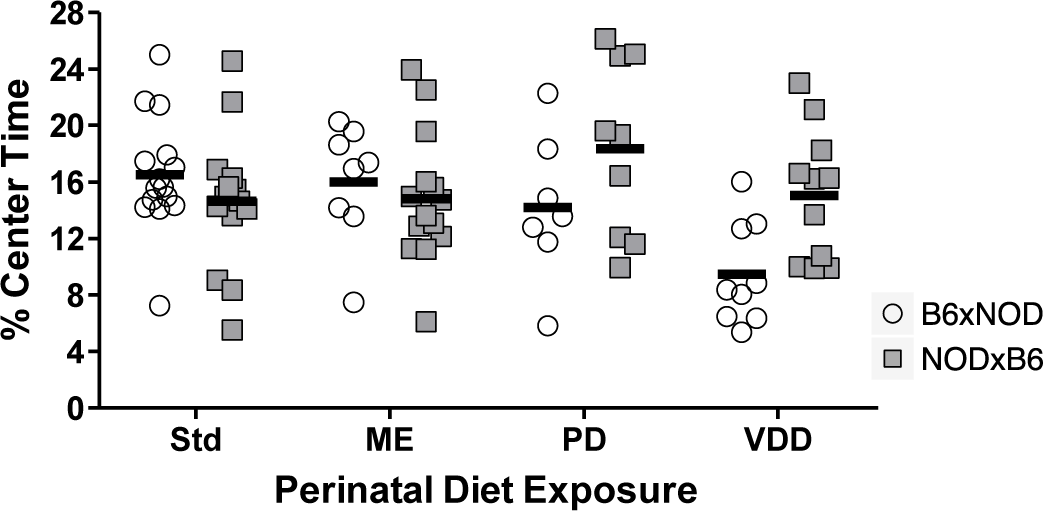
Perinatal diet-by-POE on percent center time in the 10 min OF test, for B6xNOD and NODxB6 mice exposed to Std (n=15,14), ME (n=8,14), PD (n=7,9) or VDD (n=9,11) diets; although no individual contrast is significant, diet-by-POE (p=0.0144) is significant overall.

### Effects on whole-brain gene expression

Gene expression at each microarray probeset was tested for POE, diet effects, and diet-by-POE. Significance was assessed in two ways: using the false discovery rate (FDR), and using a more conservative, permutation-based family wise error rate (FWER) threshold. The FDR (q-value = 0.05) and FWER (adjusted p-value = 0.05) thresholds were nearly identical for POE, were similar for diet-by-POE, but were over two orders of magnitude different for diet, with FWER more conservative.

#### POE detected on 15 genes, 9 imprinted

POE was FWER-significant for 15 genes (Table S9; Figure 6), a significant subset of which (nine) were imprinted (p< 2.2×10^−16^). Across the 16 genes, greater expression was not associated with either cross direction (seven more expressed in NOD×B6; ten more expressed in B6xNOD). Both patterns were seen in imprinted genes *Snord 113* and *Snord 115*, depending on the subregion (Table S9). Significant POEs clustered on (Figure 6A) chromosome 7 in the vicinity of the imprinted *Snord 115/116* family, and on chromosome 12 near the imprinted *Snord 113* family.

**Figure 6.**
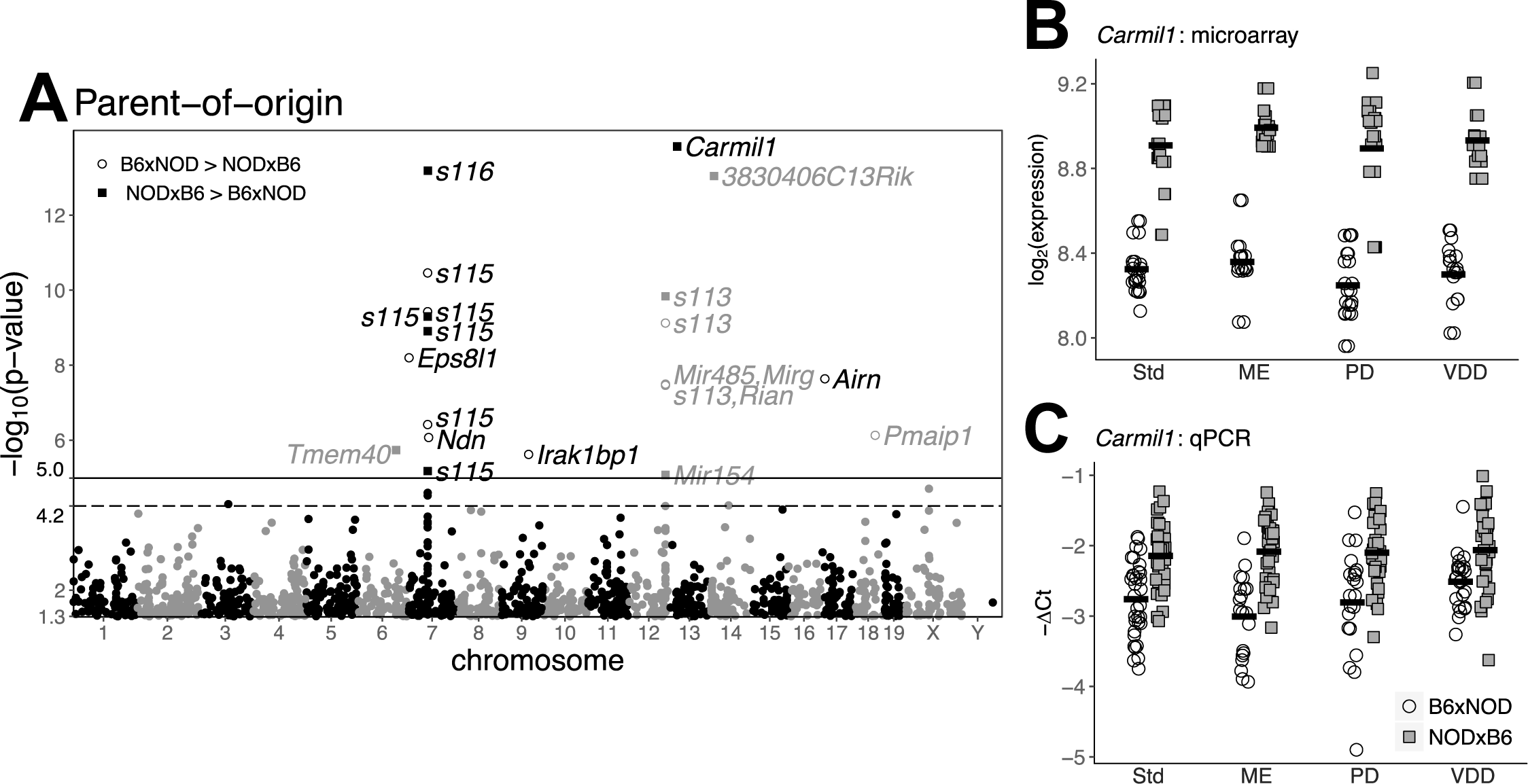
POEs on *Carmil1* gene expression. (A) Manhattan-like plot of p-values of POE on microarray-based gene expression; each point corresponds to a probeset’s genomic location, coupled with the p-value of POE on expression at that location. Probesets with a nominal p-value >.05 are not shown. The dashed and solid lines represent the FDR and FWER thresholds, respectively. Probesets above the FWER threshold are labeled with the gene(s) that they interrogate. The *S113, S115*, and *S116* labels are shorthand for *Snord 113, Snord 115, Snord 116* respectively. Labeled points are shaped according to whether expression was greater in B6xNOD or NODxB6. The most significant POE is on *Carmil1.* (B) Raw microarray expression data for *Carmil1*; circles and squares represent expression for B6xNOD and NODxB6 hybrids, respectively. POE on expression is evident under all dietary exposures. (C) qPCR validation data for *Carmil1*, showing the same significant pattern of POE in all dietary exposures, confirming the microarray findings. In any qPCR assay, increased expression *reduces* ΔCt; consequently, we use the y-axis to depict — ΔCt, ensuring that an increased y-value represents increased expression in both (B) and (C).

#### POE on non-imprinted *Carmil1* validated by qPCR

The most significant POE was on *Carmil1* (–log_10_(p) =13.8). This POE was consistent across diets (Figure 6B), and was validated by qPCR. qPCR was performed on 115 mice, 85 of which had already been assayed by microarray. POE on *Carmil1* was significant whether considering qPCR data from all 115 (p=6.3e-7), only the 85 (p=4.4e-07), or the qPCR-only 30 (p=9.7e-11) (see Table S4).

#### According to FWER, diet affects 37 (solely non-imprinted) genes

The most significantly affected was *Cnot2* (–log_10_(p) = 7.4). For 35 of the 37 genes (Table S10, Figure S5), significance was driven by the ME diet: across the 4 diets, these 35 genes were either most or least expressed in ME mice (See the “ME group rank” field in Table S10; Figure S5). By the less stringent FDR threshold, diet significantly affected 958 genes. This included even Y chromosome genes (Table S10), suggesting, since we only use females, that the FWER threshold is more appropriate.

#### According to FWER, diet-by-POE affects 16 genes, with only *Mir341* imprinted

Not only was *Mir341* the only significantly affected imprinted gene, but it was also the most significantly affected (–log_10_(p) =6.5; Table S11). However, despite *Mir341* being expected to regulate hundreds of genes (Targetscan) the 149 *(FDR selected-genes* significantly subject to diet-by-POE were not enriched for *Mir341’s* predicted regulatory targets (p=.999; using miRHub; Baran-Gale *et al.* (2013)). Following *Mir341*, the imprinted gene *Meg3* was the next most significant imprinted gene, subject diet-by-POE (but only by FDR; –log_10_(p) =4.4; Figure 7; Figure S4A). However, this weakly significant effect on *Meg3* was not reproduced in qPCR validation (Table S4).

**Figure 7.**
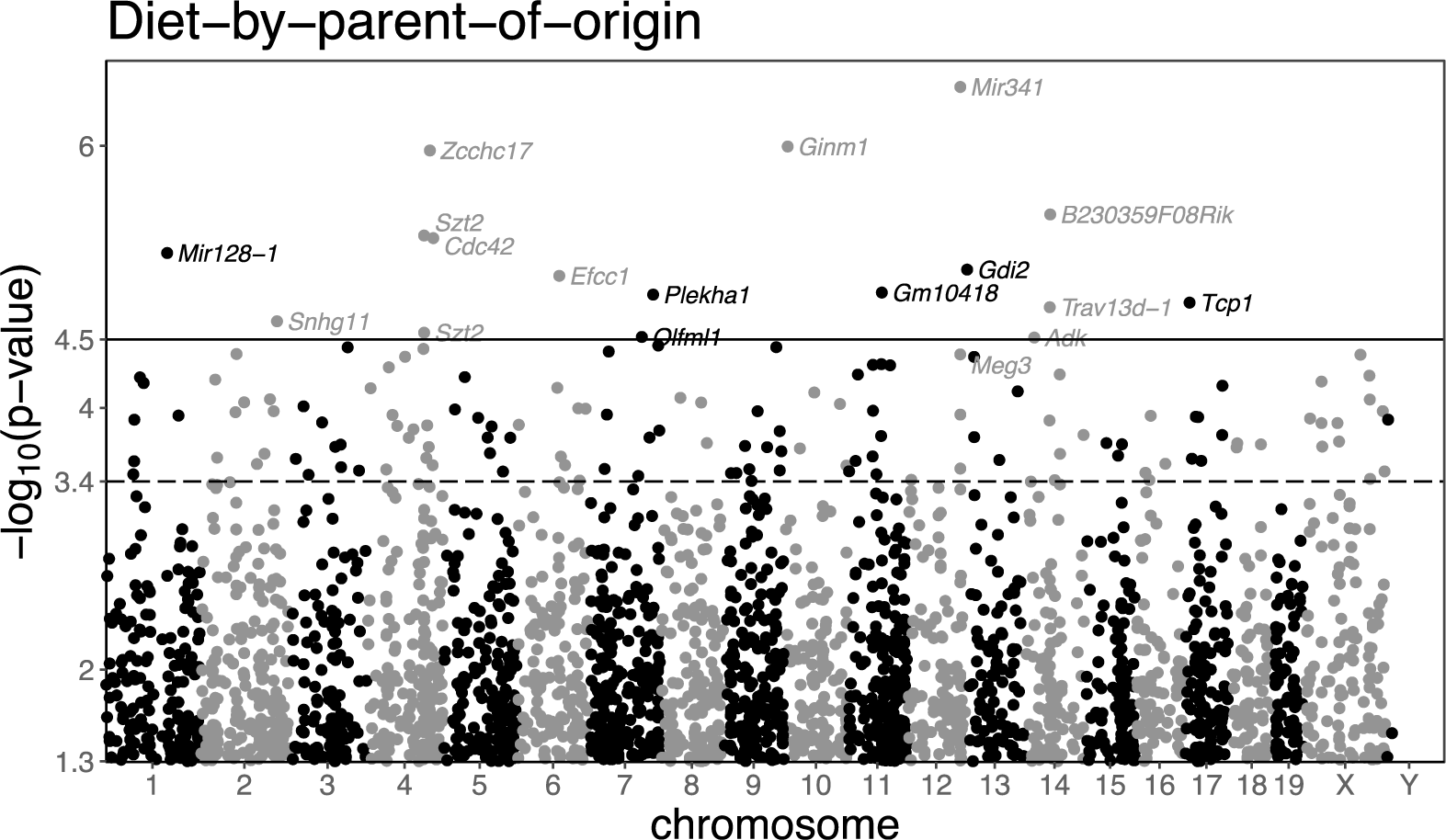
Manhattan-like plot of P-values of diet-by-POE effects on gene expression. Plotting format is similar to that used in Figure 6A. The dashed line represents the FDR threshold, and the solid line represents the FWER threshold. *Mir341* expression is the most significantly affected by diet-by-POE. Note that *Meg3*, an imprinted gene just below the FWER threshhold, is also labelled.

### Mediation of POE by way of gene expression

#### POE on the gene expression of non-imprinted gene *Carmil1* may be mediated by *Airn*

The microarray and qPCR-based evidence for POE on *Carmil1* expression raised the question: given that *Carmil1* is not known to be imprinted, might *Carmil1* expression be regulated *(i.e.*, mediated) by some imprinted gene’s expression?

We first attempted to answer this question through a ChIPBase-driven analysis (Yang *et al.* 2013) of predicted and recorded transcription factor binding sites. We found that the protein product of *Wt1*, an imprinted gene, might bind upstream of *Carmil1*— suggesting that the POE on *Carmil1* might be mediated by *Wt1.* However, we deemed this hypothesis unlikely given that, in our data, *Wt1* expression levels were unaffected by POE (p=.267).

This focused bioinformatic analysis having failed to clearly identify a mediator, we applied a genome-wide analysis: for every microarray-measured gene, we tested whether its expression mediated the POE on *Carmil1* expression. The model used to test for mediation is shown in Figure 8.

**Figure 8.**
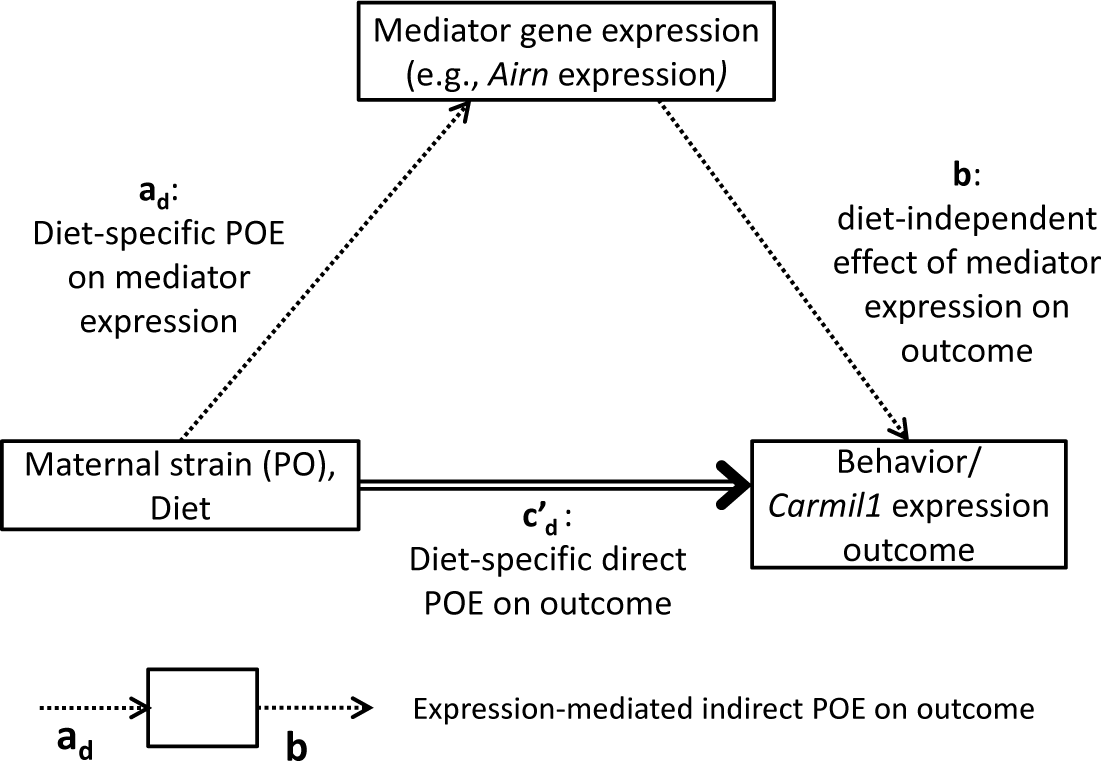
Model of gene-expression mediation of POE on the outcome, which is either behavior or *Carmil1* expression. Parent-of-origin, encoded as the maternal strain, in conjunction with diet, acts both directly upon the outcome, with effect size 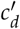, and indirectly upon the outcome, with effect size *a*_*d*_*b.* This indirect effect is composed of the diet specific POE on some mediator’s expression *(a*_*d*_) and the diet-independent effect of the mediator’s expression on the outcome (b). Not shown in this figure for clarity, but present in the actual model, are nuisance effects of dam, pipeline, batch, and behavior specific covariates, that all can affect both mediator expression and the outcome. Mediation is determined by testing whether the average indirect effect 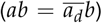 is significant according to its Combined Tail Probability (CTP).

The expression of 8 different genes was found to significantly (Combined Tail Probability, CTP<.05) mediate POE on *Carmil1* expression. For 7 of these 8 genes, their mediation *(i.e.*, indirect) effect acted against the direct effect (Figure 8); rather than explaining POE, expression of these 7 genes actually suppressed the overall POE on *Carmil1. 3830406C13Rik*, a non-imprinted protein coding gene of unknown function (Yue *et al.* 2014), was the most significant (CTP=.00289) overall mediator of POE on *Carmil1. Airn* was the most significant (CTP=.0134) mediator that was imprinted; specifically, *Airn* acted to suppress POE on *Carmil1* (Table S6; Figure 9).

**Figure 9.**
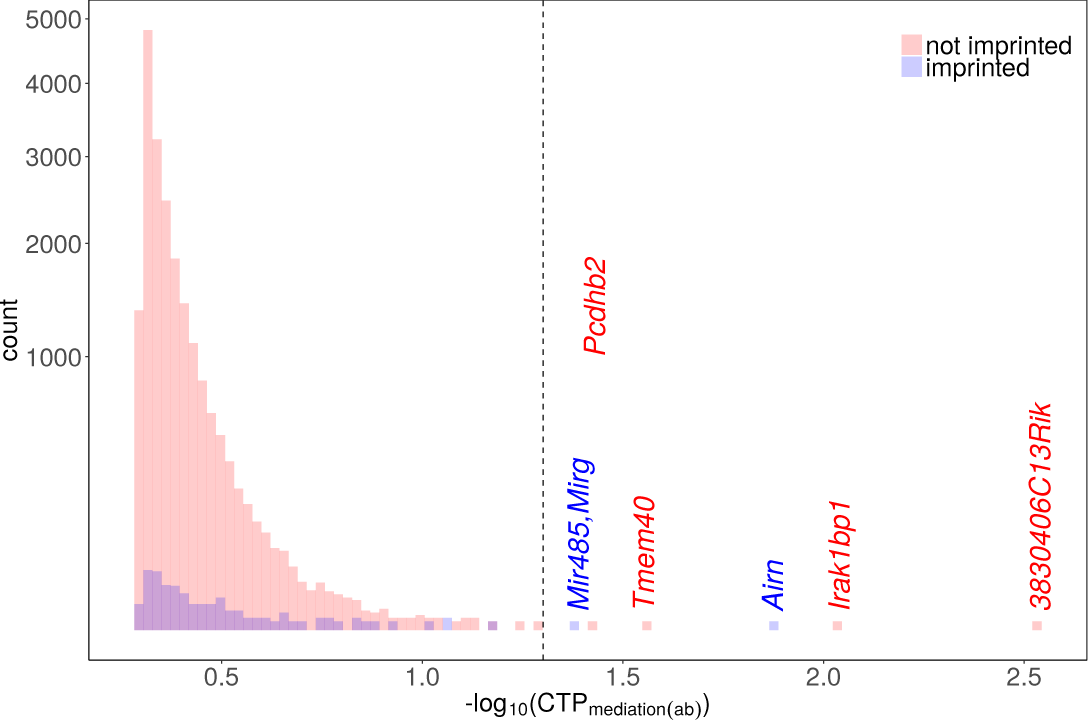
Histograms of the –log_10_ Combined Tail Probabilities (CTPs) for candidate gene mediators of POE on *Carmil1* expression. The red and blue histograms correspond to CTPs for non-imprinted and imprinted candidate mediators, respectively. Mediators whose mediation effect has a CTP<.05 (the dashed line threshold) are labeled. Notably, the imprinted gene *Airn* is one of the top 3 mediators of POE on *Carmil1.*

#### POE on behavior may be mediated by *Carmil1* and *Airn*

We repeated a similar genome-wide POE-mediation analysis for every behavioral outcome (including behaviors without significant POE in mediation-free analysis). A significant (CTP<.05) gene mediator of POE was observed for 10 of the 34 modeled behavioral outcomes. POE on some outcomes was mediated by more than one gene, and some genes mediated POE on more than one outcome. Although 16 different significant mediator-outcome pairs were observed, there were only 6 distinct genes/gene families significantly mediating any behavioral outcome: *Snord 113, Snord 115, Snord 116, 3830406C13Rik, Rian, Carmil1.* In 15 of the 16 significant mediator-outcome pairings, the gene expression mediator suppressed POE; *i.e.*, 15 of 16 gene mediators acted in the opposite direction of the direct POE on behavior (Table S7).

To determine each gene’s mediation of POE on behavior in the aggregate, we combined the CTPs for a given gene, over all behaviors, into a single metric: the “Combined Tail Probability" (CTP; Appendix J). By this metric, 21 probesets, corresponding to 17 distinct genes/gene families mediated POE on behavior in the aggregate at CTP<.05. Even though *Airn* was not a significant mediator for any individual behavior (see above), it was the most significant mediator in the aggregate (CTP=5.09e-05). *Airn* was followed closely by (a subregion of) *Snord 115* (CTP=.000408) and *Carmil1* (CTP=.000518). See Table S8.

To gain further insight into aggregate mediation, for each outcome we determined the 3 most significant POE gene-mediators. Each gene was then scored according to the number of behaviors for which it was one of the 3 top mediators. According to this metric, *Airn* was the most notable mediator, acting as one of the 3 most-significant POE-mediators for 12 behavioral outcomes, while *Carmil1* was a top-3 mediator for 8 outcomes. The enrichment for *Airn, Carmil1*, and *Snord 115* in the sets of top-3 mediators is also readily apparent in Figure S6: for each behavior, genes with a significant CTP are labelled, as are *Carmil1* and *Airn* if they were among the top 3 mediators; mediation CTPs for *Airn* and *Carmil1* are often extreme.

## DISCUSSION

Our study identifies POEs on behavior, POEs on gene expression, and shows that many of these — with notable exceptions — are robust to differences in perinatal diet. We also provide evidence for a possible explanatory pathway connecting imprinting to gene expression to behavior, and are the first study to do this.

But beyond its specific results, our study also serves to advance a general protocol based on reciprocal F1s for studying POE and perinatal environment effects on a complex trait. The RF1 holds genetic background constant while varing parent of origin, making it the most powerful design for detecting POE. To investigate the interaction of developmental-environment with POE, we further varied *in utero* nutrition, using four different diets: diet was a relatively easy variable to control, and ample evidence suggested its importance in POE. By repeating the behavioral and expression assays under multiple dietary conditions, we: 1) enabled detection of environment-by-POE, 2) hedged our bets, as an effect that would be unobservable in one environment might be amplified in another, and 3) enabled detection of POE that generalizes across environments.

In the remainder, we discuss the range of mechanisms that might explain POE as discoverable by our approach; our specific results on POE, diet, and diet-by-POE; and lastly, we reflect on the use of replicable vs non-replicable populations for POE discovery and investigation.

### Coding-POE vs eQTL-POE, and POE observability

The two examined groups of female RF1s, NOD×B6 and B6×NOD, were (aside from mitochondria) genetically identical. Consequently, differences in phenotype between these two groups could with high probability be attributed to POE. But for such observable POE to exist, imprinting/maternal factors must have interacted with a locus differing in sequence between parents (Figure 10A). This difference driving the POE could have been in a coding region, making it a “coding-POE”, and/or in a regulatory region, making it an “eQTL-POE”.

**Figure 10** Examples of cis/trans coding-POE and eQTL-POE, in RF1s. Examples depict an imprinted gene which is fully active when maternally inherited, but fully silenced when paternally inherited. Similar examples could be constructed for maternal effects. (A) A lack of observable POE in spite of imprinting: B6 and NOD are identical in sequence, so whichever allele is silenced, the resulting expression product is the same in both RF1 directions. (B) Coding-POE: B6 and NOD differ in coding sequence causing allele-specific expression differences between RF1 directions (unobservable by microarray). (C) eQTL POE: the NOD promoter attracts a more effective transcription factor (TF), so NODxB6, in in which the NOD allele is expressed, yields more expression. (D) eQTL-POE driven by background-dependent imprinting: imprinting is lost in B6, so NODxB6, in which imprint-silencing affects neither allele, yields more expression. (E) Trans coding-POE upon locus 2: locus 2 is identical between NOD and B6, but it is regulated by an imprinted TF whose NOD version is more efficient; so in NODxB6, in which the NOD TF is expressed, locus 2 expression is increased. (F) Trans eQTL-POE upon locus 2: locus 2 is regulated by an imprinted TF whose NOD version has a stronger promoter; so in NODxB6, in which the NOD allele for the TF is expressed, increased availability of the TF increases microarray-observable locus 2 expression.

In coding-POE, the expressed allele’s coding sequence differs between the two cross directions. Consequently, the RF1 populations are equal in total expression, but allele-specific expression (ASE) differs (Figure 10B). Although the microarrays in our study cannot quantify ASE, ASE differences can still manifest as an observable POE on an emergent phenotype such as behavior, or as POE on total expression of a downstream gene.

By contrast, eQTL-POE could arise by way of non-coding cis-eQTLs that alter total expression of an imprinted gene. For example, an eQTL-POE could arise from differences in promoter attractiveness, (Figure 10C). Or, perhaps more interestingly, eQTL- POE could arise by way of genetic background-dependent loss of imprinting (Vrana 2007; Duselis *et al.* 2005; Wolf *et al.* 2014) (Figure 10D). In our study, all directly-observed POE on expression are necessarily instances of eQTL-POE, because we did not employ assays capable of measuring ASE.

eQTL-POE and coding-POE both require a genetic difference between parents in some imprinted or maternally-affected gene. However, any gene can exhibit POE— provided it is regulated by the imprinted/maternal-effect gene. This trans effect can occur eitherby way of coding-POE (Figure 10E) or eQTL-POE (Figure 10F).

Both types of POE may be undetected by our study. As mentioned above, coding-POE is unobservable by our microarrays. Additionally, by measuring expression once, ~8 weeks after birth, we may have failed to observe POE during transient, developmental-stage-specific imprintin. And by measuring whole-brain gene expression, we may have occluded POE arising from imprinting that is specific to subregions of the brain (Koerner *et al.* 2009; Prickett and Oakey 2012).

### POE on expression

All 9 imprinted genes that were subject to POE contain non-coding variants that differ between NOD and B6, a finding consistent with cis-driven, eQTL-POE (Figure 10C,D). However, six of the genes subject to POE were non-imprinted, including *Carmil1.* POE on such genes may be driven by maternal effects, or perhaps by trans-acting imprinted regulators (as in Figure 10F).

### Mediation of POE on *Carmil1*

To determine potential imprinted regulators of *Carmil1* expression, we applied mediation analysis, identifying *Airn.* Unexpectedly however, *Airn* exerted its mediation effect in the opposite direction of the overall POE on *Carmil1 (ab* and *c′* have opposite signs in Table S6), suggesting that *Airn* suppresses POE on *Carmil1* in trans. All but one of the other significant mediators also acted as POE suppressors. The lack of explanatory mediation in the same direction as the overall POE may be due to the many unobservable forms of POE on expression: genes that fail to exhibit POE in their own expression cannot be statistically significant mediators of POE on another gene’s expression. Alternatively, *Airn* and the other imprinted mediator may be suppressing unobserved maternal effects on *Carmil1.*

### POE on behavior and its mediation by gene expression

Five behaviors were significantly affected by POE, four of which were locomotor behaviors. The enrichment for POE could in part be due to increased power: locomotor activity has been found to be among the most stable of behaviors across laboratories and time (Crabbe *et al.* 1999; Wahlsten *et al.* 2006), resulting in more power to observe group differences. Also, however, given that locomotor activity has been used to measure rodent emotionality (Hall 1934) and predict addiction-related behavior (Piazza et *al.* 1989), our POE results on locomotor activity may suggest that POEs are in fact important determinants of emotionality and/or addiction.

For the 5 POE-significant behaviors (and the other non-POE-significant behaviors too), POE must have been driven by some gene subject to POE; to identify such genes, we applied mediation analysis, finding 17 genes that mediate behavioral POE. However, for 16 of the 17 genes, the estimated mediation effect was to suppress POE; i.e, these genes did not explain the overall POE on behavior. We posit that explanatory POE on gene expression may simply have been unobservable, for the reasons described earlier.

### *Airn* and *Carmil1* as mediators of POE

The most commonly shared mediators of behavior were *Carmil1* and *Airn*, with *Airn* also being the top mediator of POE on *Carmil1.*

*Airn’s* mediation of POE is likely trans-acting. *Airn* is an imprinted, paternally-expressed, long non-coding RNA (lncRNA), which to our knowledge has not been found to affect any complex trait directly. Rather, *Airn* is known to control imprinting of three nearby maternally-expressed genes: *Slc22a2, Slc22a3*, and *Igf2r* (Cleaton *et al.* 2014). But none of the three genes were at all significant mediators of POE on any outcome of interest in our dataset. So, akin to other lncRNAs and imprinted genes found to affect distal gene expression (Vance and Ponting 2014; Gabory *et al.* 2009), we posit that *Airn* may be exerting POE on behavior by affecting distal genes, such as *Carmil1* or *Snord 115* (as in Figure 10E). Our study is underpowered to directly examine this two-step mediation hypothesis.

*Carmil1* may provide a link between cytoskeleton dynamics and cell migration, and behavioral change. *Carmil1* has a known cellular role in: 1) interacting with Capping Protein, which regulates actin elongation; and 2) activating the small GTPase *Rac1*, an important regulator of cytoskeletal dynamics (Gonzalez-Billault *et al.* 2012). Such actin cytoskeleton dynamics, critical for cytokinesis and cell migration (Rottner *et al.* 2017), are important throughout the lifespan for neurodevelopment and neural plasticity (Menon and Gupton 2016; Gordon-Weeks and Fournier 2014). In *C. elegans*, neuronal cell and axon growth cone migration has been shown to be negatively regulated by *CRML-1*, the homolog of *Carmil1* (Vanderzalm *et al.* 2009). Our study, in a mammal, is the first to find a direct association between variation in *Carmil1* expression and behavior.

### Caveats to mediation analysis of POE on *Carmil1* and behavior

We note that our analysis was applied one candidate mediator at a time; thus, any significant mediators may simply be co-expressed with the true mediator gene(s). We also note that for both mediation analyses (*Carmil1*/behavior outcome) we assumed a direction of causality in which some imprinted gene mediates POE on the outcome; although this might seem intuitive, it cannot be verified, and the “outcome” might actually mediate the imprinted gene.

Our directionality assumption is particularly uncertain in the behavioral analysis: expression in the brain was, out of necessity, measured after behavior; consequently, stressful behavioral assays could have altered expression. In future studies, we intend to address this weakness by a matching-based imputation: behavior-unperturbed expression will be imputed in behaviorally-assayed mice using expression data from mice that were unexposed but are genetically identical and otherwise perfectly matched (cf. related matching-based designs in Crowley *et al.* 2014)

### Diet effects

Our data revealed significant diet effects on gene expression, with significance primarily driven by extremely low/high expression under the ME diet. This may be unsurprising given the direct role of methyl donors on DNA methylation and, consequently, on the regulation of gene expression. Future perinatal-diet studies may benefit from a ladder of methyl enrichment values.

Notably, diet significantly altered the expression of only 37 genes according to the strict FWER threshold, but 958 according to the FDR threshold. Some of these additional hits are likely false-positives (e.g. Y chromosome genes). But it is conceivable that diet did in fact cause a systemic, diet-buffering change to the overall network of gene expression levels (MacNeil and Walhout. Indeed, our FDR numbers seem consistent with earlier work: in the few examples (to our knowledge) of FDR corrected results from previous rodent studies of perinatal diet effects on gene expression (Mortensen *et al.* 2009; Altobelli *et al.* 2013; Barnett *et al.* 2015), 500-1000 genes were differentially expressed.

Although diet significantly altered the expression of numerous genes, the only complex phenotypes affected were body weight and PPI, and those effects were barely significant. The lack of significant diet effects on behavior, even in the presence of expression changes, is surprising but not entirely unexpected. Among other possibilities, the diet effects on behavior may be too small to overcome a sample size that was split among four different diets. Moreover, we measured a limited set of behaviors that may not have been altered by diet-driven gene expression changes.

### Diet-by-parent-of-origin effects

Although our study perturbed nutrients involved in imprinting, the only imprinted gene subject to diet-by-POE and passing FWER was *Mir341.* The next most significant imprinted gene, which passed FDR but not FWER, was *Meg3*. However, these results are both uncertain: the genes predicted to be regulated by *Mir341* do not seem to manifest diet-by-POE effects in our data; and *Meg3’s* diet-by-POE was observed in microarray data but failed to replicate in qPCR data (Figure S4).

In addition to inevitable lower power for testing interaction effects, the relative lack of observed diet-by-POE on imprinted genes may also be in part due to the aforementioned transience and/or tissue specificity of some imprinted genes (Ivanova *et al.*, or because our diets are insufficiently extreme to elicit diet-by-POE. Insufficiently extreme diets may also contribute to lack of diet-by-POE on most of our behaviors (save for percent center time).

As for the 16 non-imprinted genes subject to diet-by-POE (by FWER), these may be regulated by imprinted genes that are subject to the aforementioned unobservable diet-by-POE (Figure 10E,F). Or, perhaps more likely, the 16 imprinted genes are controlled by maternal effects.

### Studying POE in replicable vs non-replicable (outbred) populations

A number of previous studies of POE on complex traits have used outbred populations, such as F2, backcross, or heterogeneous stocks (Lawson *et al.* 2013). The advantages of such outbred populations over the RF1 are that: 1) POE can be detected simultaneously with non-POE genetic effects; and 2) POE arising from imprinting vs maternal effects can be disambiguated— a significant difference between reciprocal heterozygote (at some locus) offspring from heterozygote (at that locus) mothers can be ascribed to imprinting rather than to a maternal effect (Hager *et al.* 2008).

However, outbred populations have disadvantages as well: due to the fact that every animal is genetically distinct in an outbred population, alternate parent-of-origin states can never be observed in the exact same genetic background; this confounding limits the power of outbred populations to estimate POE. By contrast, in the RF1, individuals of alternate parent-of-origin state can always be perfectly matched in the same genetic background (save for the mitochondrial genome), allowing unconfounded and unbiased estimates of the causal POE. Moreover, whereas the irreplicability of outbred animals makes it impossible to perfectly recreate genetic state for a validation study, *e.g.*, a future study evaluating the effect of a knockout of *Carmil1* on behavioral POE, this is readily available for the RF1.

Only a handful of other studies have used an RF1 strategy to study POE on complex mammalian traits such as behavior. But none of these studies (save for a recent one of our own in Schoenrock *et al.* (2017)) simultaneously varied environment. Nor have other RF1 studies simultaneously measured gene expression. We are the first to demonstrate that combining the RF1 design, environmental perturbation, and observation of gene expression, provides a powerful paradigm for studying environment-by-POE on a complex mammalian trait.

## ACKNOWLEDGEMENTS

This work was funded by a National Institute of Mental Health (NIMH)/National Human Genome Research Institute Center of Excellence for Genome Sciences grant (P50-MH090338 and P50- HG006582) FPMdV, and by NIMH R01-MH100241 to WV and LMT. DO also received partial support from a PhRMA Foundation predoctoral informatics fellowship. Microarray studies were performed in the UNC functional Genomics Core under the direction of Mike Vernon. miRHub guidance was provided by Matt Kanke and Sara Selitsky.

## APPENDIX A: BEHAVIOR MODELS

The LMM used to model behavioral phenotypes (excluding the startle/PPI phenotypes) was as follows. The behavioral outcome *y*_*mi*_ of mouse *mi* was modeled as

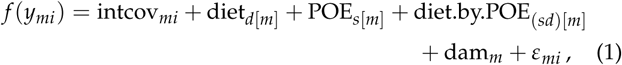

where *mi* denotes the *i*th mouse of mother *m*; *d*[*m*] denotes mother *m*’s diet, where *d* = 1, …, 4, corresponding to diets Std, ME, VDD and PD; *s*[*m*] denotes the mother’s strain, where *s =* 1,2 corresponds to B6 and NOD respectively; (*sd*)[*m*] denotes the mother’s diet and strain combination. Modeled effects consisted of: intcov_*mi*_, a fixed intercept and a set of (behavior-specific) fixed effect covariates; diet_*d*_, a fixed effect of diet_*d*_; POE_*s*_, a fixed effect of POE (technically, strain-by-POE); diet.by.POE_*sd*_ a fixed effect of diet-by-POE; and dam_m_, a random effect of dam. The function *f* () is a transformation chosen to ensure the residuals *ε*_*mi*_ are approximately normal (Appendix B).

### Startle/PPI Models

For every prepulse intensity, 6 measurements of the average startle response were taken per mouse (all in the same chamber). The startle/PPI LMMs therefore accounted for repeated measures. Letting *y*_*mi*,*j*_ be mouse *mi*’s *j*th measurement, we modeled:

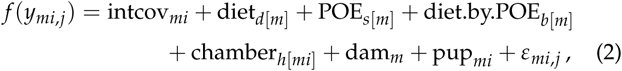

where chamber_*h*[*mi*]_ is a random effect of chamber, and pup_*mi*_ is the random effect of mouse *mi*.

## APPENDIX B: VARIABLE TRANSFORMATION PROCEDURE

A transformation procedure was applied to both the expression and the behavior phenotypes to ensure residual normality. For a given LMM requiring a transformation of the outcome *y*, the procedure was as follows. Center and scale *y* to mean 0 and standard deviation 1 to give z. Apply a shifted Box-Cox transformation (Sakia 1992; Box and Cox 1964), restricted to the ladder of powers λ ∈{–3, –2, –1, –.5,0,.5,1,2,3} to give in each case values *z*^(λ)^. For each transformation *z*^(λ)^, the LMM is fitted, and residual normality is evaluated using the Shapiro-Wilk *W* statistic (Shapiro and Wilk 1965); denote the optimal λ as, 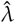. If 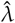 ∈ {0,.5,1,2}, then use 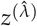; if 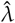 ∈ {–2, –1, –.5}, then additionally negate the value, in order to ensure the monotonicity of the transformation and thereby improve interpretability of effect estimates; if the 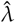 *∈* {–3,3}, then discard the transformation and instead apply a rank inverse normal transform (Van der Waerden 1952). Rescale the selected transformed variable to mean 0 and standard deviation 1.

## APPENDIX C: MICROARRAY EXPRESSION MODELS

Expression was first adjusted by regressing out nuisance factors, and then the *adjuste*d expression was modeled to test diet, POE, and diet-by-POE. This two-step process was employed to facilitate permutation testing later on.

### Generation of the adjusted expression outcome

Letting *y*_*mi,j*_ be the average expression of probes in probeset *j* for mouse *mi*, we obtained adjusted expression values as residuals 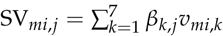_*mi,j*_from the linear model:

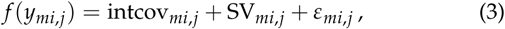

where the covariates in intcov_*mi,j*_ were the nuisance effects of pipeline and behavioral batch. The SV_*mi,j*_ term modeled fixed effects for 7 “surrogate variables” (SVs), which represented aggregate effects of unobserved confounding on the microarray (Appendix F). Specifically, 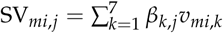, where 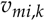 is mouse *mi*’*s* value for the *k*th SV, and *β*_*k,j*_ is the fixed effect of that SV on the expression of probeset *j*. (Estimation of the SVs themselves is described in Appendix D.)

### Model of adjusted expression outcome

For each probeset *j*, adjusted expression (a.k.a., the residuals from Eq 3) was then analyzed using the LMM,

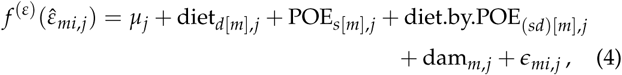

where *μ*_*j*_ and ∊_*mi,j*_ are the intercept and residual error, *f*^(*ε*)^ is a transformation that may be different from *f* in Eq 3, and other terms are defined as in Eq 1.

## APPENDIX D: SURROGATE VARIABLE ESTIMATION ALLOWING FOR RANDOM EFFECTS

Gene expression measurements by microarray are typically affected by many unobserved factors, some of which can have a large confounding effect on transcript levels across many genes. One way to control for such unobserved factors is to first model their aggregate effects as linear combinations of “surrogate variables” (SVs; Leek and Storey 2007), and then include these SVs as predictors in subsequent modeling, and/or regress these effects out (as in Appendix C).

Here we mostly— deviating somewhat to accommodate random effects and variable transformation— follow the Supervised Surrogate Variable Analysis (SSVA) approach of Leek (2014), which defines the SVs using negative control probes; success of this approach requires that unobserved confounding effects arise from technical rather than biological variation. As a further aside, we note that our approach is also largely equivalent to the “remove unwanted variation with negative control genes” (RUVg) strategy (Risso *et al.* 2014), applied to microarray data.

In our implementation of SSVA, we first estimate a standardized matrix of the aggregate effects that arise from unobserved factors, *E* For each negative control probe *c =* 1,…, C, we fitted the LMM

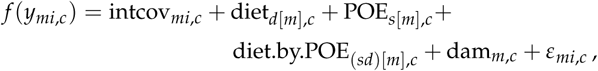

where terms are defined as in Eq 1 and Eq 4, and where the estimated residuals, 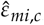, were standardized and stored in n-vector *e*_*c*_. These steps were repeated for all *C* negative control probes to give the *n* × *C* matrix **E**.

Let the SVD of *E* be denoted as **UΣV′**. Under this parameterization, the space of aggregate unobserved factor effects on the control probes is (by construction) spanned by the *n* columns of **U**. Since a model for main probes that included all *n* columns as surrogate variables would be unidentifiable, the first *K =* 7 columns of **U** were chosen as an approximating subset of surrogate variables. *K =* 7 was chosen by following the strategy described in Sun *et al.* (2012) for K-selection in SVA with random effects: a plot of the squared eigenvalues from **Σ** was examined, and it revealed an inflection point at 7 eigenvalues.

Of note, the original implementation of SSVA did not regress any effects out of control probes, under the assumption that these probes should be unaffected; in contrast, we regress these effects out before computing eignevectors. We justify this by noting that if in fact the treatments of interest somehow did affect the control probes, we would not want these treatment effects to be incorporated into the surrogate variables. And if the control probes truly are unaffected by any of the observed experimental factors, then there should be no harm in residualizing out these size-zero effects.

## APPENDIX E: BIAS-ADJUSTMENT FOR GENE EXPRESSION P-VALUES

For some effect types that were tested in the gene expression model of Eq 4, the distribution of nominal p-values across all transcripts was consistent with those p-values being downwardly biased. To remove this bias, which would otherwise invalidate our use of FDR, we applied an empirical adjustment similar to the genomic control procedure of Devlin *et al.* (2001) (see also Dadd *et al.* 2009). Let *pj* be the p-value associated with a given effect type (diet, POE, or diet-by-POE) on the jth probeset, let *F(x)* be the cumulative distribution function for the 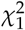 density, and define *x_j_ = F^—^*^1^ (*p*_*j*_) and ***x*** = (*x*_1_,…, *x*_*m*_). Under unbiasedness, p-values associated with testing for given effect should, under the null, have a uniform distribution, *p*_*j*_ ~ Unif(0,1), such that *x*_*j*_ ~ 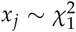. Assuming most results are in fact null, in the dataset as a whole we would expect median(**x**) ≃ *F*^*—*1^(0.5). However, if significances were systematically inflated, the null 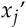’s would appear as if from a scaled 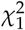 such that 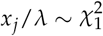 with inflation factor λ > 1. Therefore, we correct for this systematic inflation by first estimating the inflation factor as 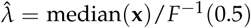 and then calculating bias-adjusted p-values as 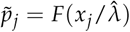.

## APPENDIX F: PERMUTATION-BASED FWER THRESHOLDS FOR GENE EXPRESSION P-VALUES

For gene expression, empirical p-value thresholds that controlled for the family-wise error rate (FWER) across all probesets were determined by permutation. A separate FWER threshold was computed per effect of interest (diet, parent-of-origin, and diet-by-parent-of-origin). Below, we describe the permutations that were generated, the statistic that was collected per permutation, and how this was translated into a significance threshold.

### Structure of permutation

For every permutation-tested effect type, we generated a separate set of *W =* 401 permutations (including the identity permutation), *w =* 1…, W. Litters were taken as exchangeable units; diet/strain labels were permuted amongst the dams, and all pups of a given dam were assigned their dam’s diet/strain label. Permuting *labels*, rather than outcomes, enabled us to allow for varying litter sizes between dams.

For the main effects we employed a form of restricted permutation (Anderson and Braak 2003; Good 2005); i.e., for parent-of-origin effects, we randomly permuted the strain labels (s in Eq 4) between dams that had been *exposed to the same diet*, whereas for diet effects, we randomly permuted diet labels (**d**) between dams *of the same strain.*

For the interaction effect of diet-by-POE, we employed a form of unrestricted permutation (Anderson and Braak 2003; Good 2005) of the interaction labels. In particular, we permuted the interaction labels **g** between dams. However, the **s** and **d** labels were held constant even as the interaction labels **g** were permuted.

### Permutation statistic and threshold computation

For each permutation *w* and probeset *j =* 1,…, *J* we fitted the expression LMM of Eq 4. Note that the modeled outcome in this equation is adjusted gene expression from which *all nuisance co-variates have already been regressed*; following Gail *et al.* (1988), this residualization was performed to facilitate exchangeability for the effects of interest. For every permutation, the fitting of 4 included recalculation of the transformation *f*^(ε)^. Furthermore, for every permutation, we bias-adjusted (through genomic control, Appendix E) the p-values, 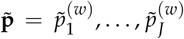 and recorded the minimum, *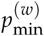*.

The set of *W* such minimum p-values from all permutations was then used to estimate the FWER *α =* 0.05 threshold via modeling of a generalized extreme value (GEV) distribution after Dudbridge and Koeleman (2004); Manly (2006). Specifically, a GEV was fitted to 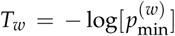 for *w =* 1,…, *W* using R package evir (Pfaff and McNeil 2012), and the fitted GEV was used to estimate the upper 5% quantile, *T*_*α*_=.05. *T*_*α*_=.05 was then translated back into a threshold on the p-value scale as *p*_*α*_=.05 = *e*^−*Tα*=.05^. Note that, as a conservative measure, the GEV fit included the identity permutation.

## APPENDIX G: PROBE ALIGNMENTS AND ESTIMATED PROBESET POSITIONS

Probe alignments were downloaded from the Ensembl 38.75 func-gen database (Yates *et al.* 2016). Notably, this database contained alignments for MoGene1.0 ST probes, rather than for the MoGene1.1 ST probes that we used in our experiment. To address this mismatch, we imputed 1.1 alignments by using the fact that every 1.1 probe is identical to at least one 1.0 probe in sequence (though not in probe id); we formed correspondences from each 1 probe to its identical-sequence 1.0 probe alignment. Since most probes aligned to multiple positions, we estimated per probe and per probeset, the “intended” target position, defining this self-referentially as the position that minimizes the sum of distances between probes in the same probeset.

## APPENDIX H: CRITERIA FOR MASKING BIASED AND UNINFORMATIVE PROBES/PROBESETS

APT masking was used to eliminate four types of probes: 1) probes aligning to ≥ 100 locations; 2) probes aligning outside of annotated exons; 3) probes whose “interior” (basepairs 3-21) aligned to regions in which NOD possesses a variant relative to B6, i.e., probes with a binding affinity difference between strains (Dannemann *et al.* 2009), where NOD variants were extracted from the Inbred Strain Variant Database (Oreper *et al.* 2017); or 4) redundant probes mapping to the same position. Following probe masking, probe*sets* were eliminated if they contained <4 non-masked probes, or if every remaining non-masked probe measured <32 units of expression across all samples.

## APPENDIX I: QPCR ANALYSIS qPCR model

Letting *y’*_*mi,j*_ be the qPCR relative cycle threshold for a targetted gene (*Meg3* or *Carmil1*), we modeled:

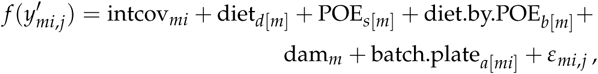

where intcov includes the intercept and behavioral pipeline, batch.plate_*a*_ is a random effect of the combination *a* of breeding batch and qPCR plate, and the other terms are akin those defined in the microarray model (Appendix C).

### qPCR normalization

The raw value measured by qPCR is a target gene’s cycle threshold. To allow comparison between qPCR batches, which can vary in replication efficiency, the cycle threshold for a target gene must be normalized by some reference gene that is unaffected by biological factors. As such, rather than modeling the cycle threshold, we model the relative cycle threshold, defined as Δ*Ct* = *Ct*_target_ — *Ct*_reference_. The Δ*Ct* relative cycle threshold represents the relative gene expression level of the target gene on the log scale (Didion *et al.* 2015). The larger Δ*Ct* is, the less the target gene expression.

We chose *Rfng* as the reference gene, because microarray data suggested negligible effects of diet, POE and diet-by-POE on *Rfng* expression. Specifically, each candidate reference gene was assigned a score equal to the minimum of the p-values for POE, diet-by-POE, and diet effects on the candidate reference’s microarray-measured expression. *Rfng* had the largest such score.

## APPENDIX J: BAYESIAN MEDIATION MODEL

Mediation analysis is typically posed as the estimation of the model in Figure 11: An intervention or predictor variable *X* affects an outcome *Y* either directly or/and through an observed mediator outcome *M*. In our case, *X* is reciprocal direction (*i.e*., parent-of-origin, coded as the maternal strain), *M* is the expression of a mediator gene, and *Y* is the outcome of primary interest, either expression of *Carmil1* or a behavioral phenotype. By common convention, the effect of *X* on M, *i.e.*, the POE on M, is denoted *a*, which in our case is *a*_*d*_ to allow different effects under each diet *d*, and the effect of *M* on *Y* is denoted *b*. The product *a*_*d*_*b* is then the expression-mediated effect of parent-of-origin on *Y*, conditional on the diet *d*, and our primary quantity of interest is this value averaged over diets, 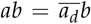. The direct effect of *X* on *Y* after accounting for mediation by *M* is denoted *c*’, which in our case is analogously diet-specific and denoted here as 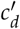 with average direct effect 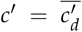. (Not explicitly calculated here but used elsewhere is *c*, which would be the effect of *X* on *Y* if mediation were unmodeled.) When *ab* and *c*^’^ have opposite signs, mediation by way of *M* acts to suppress the overall parent-of-origin effect on the outcome *Y*.

**Figure 11.**
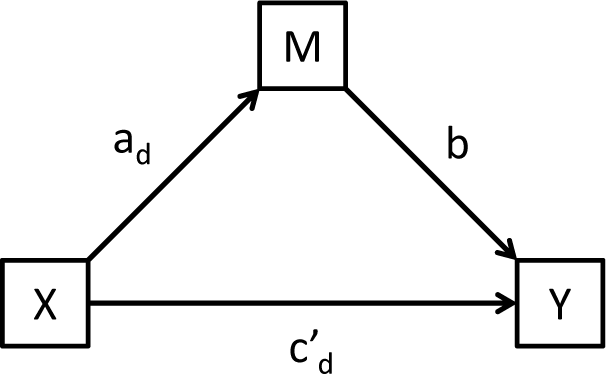
Multilevel mediation model in which the levels are diets. *X* represents the maternal-strain treatment, *Y* is the outcome (behavior or expression), and *M* is the mediating gene expression factor. *a*_*d*_ is the (diet-specific) effect of the treatment on the mediator, *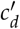* is the (diet-specific effect) direct effect of the treatment on the outcome, and *b* is the (diet-independent) effect of the mediator value on the outcome.

### Linked LMMs

Our mediation model for the effect of a gene-expression-mediator *z* on an outcome *y* is specified via two linked LMMs as:

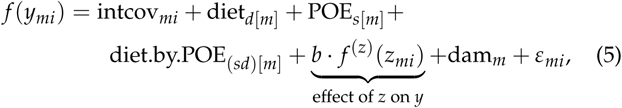

where *f*^(*Z*)^ denotes a transformation that may be different from *f, b* is the effect of mediator *z* on *y*, and the combined contribution of POE and diet.by.POE provides the direct effect *c*’_*d*_. Meanwhile, mediator *z* is simultaneously modeled as

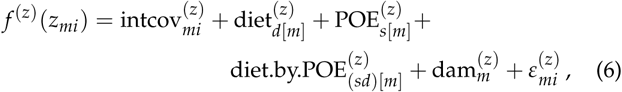

where, for example, the notation intcov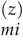 means the same regression input as intcov_*mi*_ but with regression coefficients specific to mediator *z* rather than outcome *y*, and the combined contribution of POE^(z)^ and diet.by.POE^(*z*)^ provides the effect *a*_*d*_. Specifically, the correspondence of Eq 5 and Eq 6 to the more general mediation analysis is as follows:

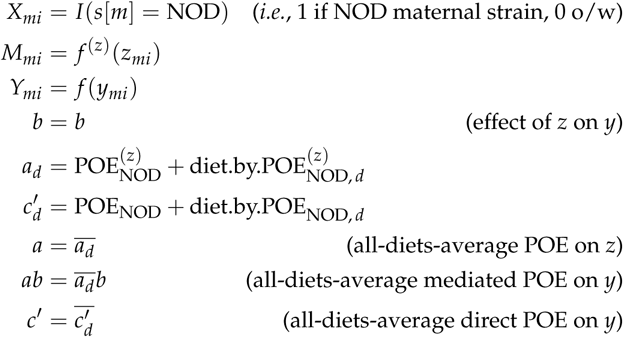

where POE_NOD_ is the effect of switching from an NOD mother to a B6 mother, and diet.by.POE_NOD,*d*_ is the additional effect of this for diet *d*.

### Transformations, expression adjustment, priors, and MCMC sampling

Prior to fitting the Bayesian mediation model, all candidate mediators and outcomes were transformed using the same process as described earlier; *i.e.*, transforms were chosen to ensure normality using the frequentist, mediation-free models (Appendix B, C). Additionally, akin to the mediation-free microarray analysis, surrogate variable effects (Appendix D) were regressed out of every gene’s expression prior to mediation modeling. However, unlike the mediation-free analysis of expression, batch and pipeline were *not* regressed out, and were included as nuisance effects on mediator and outcome in the mediation model. Priors were specified as follows, noting that *M* and *Y* by construction have means of 0 and standard deviation 1: fixed effects *(i.e.*, all effects except dam) were given priors of N(0,5^2^); and the random effect of dam was modeled as drawn from N(0, *τ*^2^) with *τ*^2^ ~ Unif(0,25). Model fitting proceded by running a single MCMC chain for 16,000 timesteps, of which the first 3,200 were discarded *(i.e.*, as burn-in), and the last 12,800 were retained for estimation.

### Combined Tail Probability: a statistic to quantify aggregate mediation

To quantify the extent to which a given gene’s expression mediated POE on multiple outcomes, we use a statistic inspired by the Fisher combined p-value that we refer to as the “Combined Tail Probability” (CTP). The CTP is the probability that a value drawn from 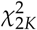 is at least as extreme as the statistic 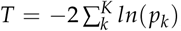, where *K* is the number of outcomes tested for mediation, and *p*_*k*_ is the CTP for the mediator’s indirect effect on outcome *k*. Although the implicit distributional assumption is not strictly justified, the CTP associated with *T* provides a statistic for evaluating which mediators are strongest in aggregate.

## SUPPLEMENTAL MATERIAL

**Figure S1.**
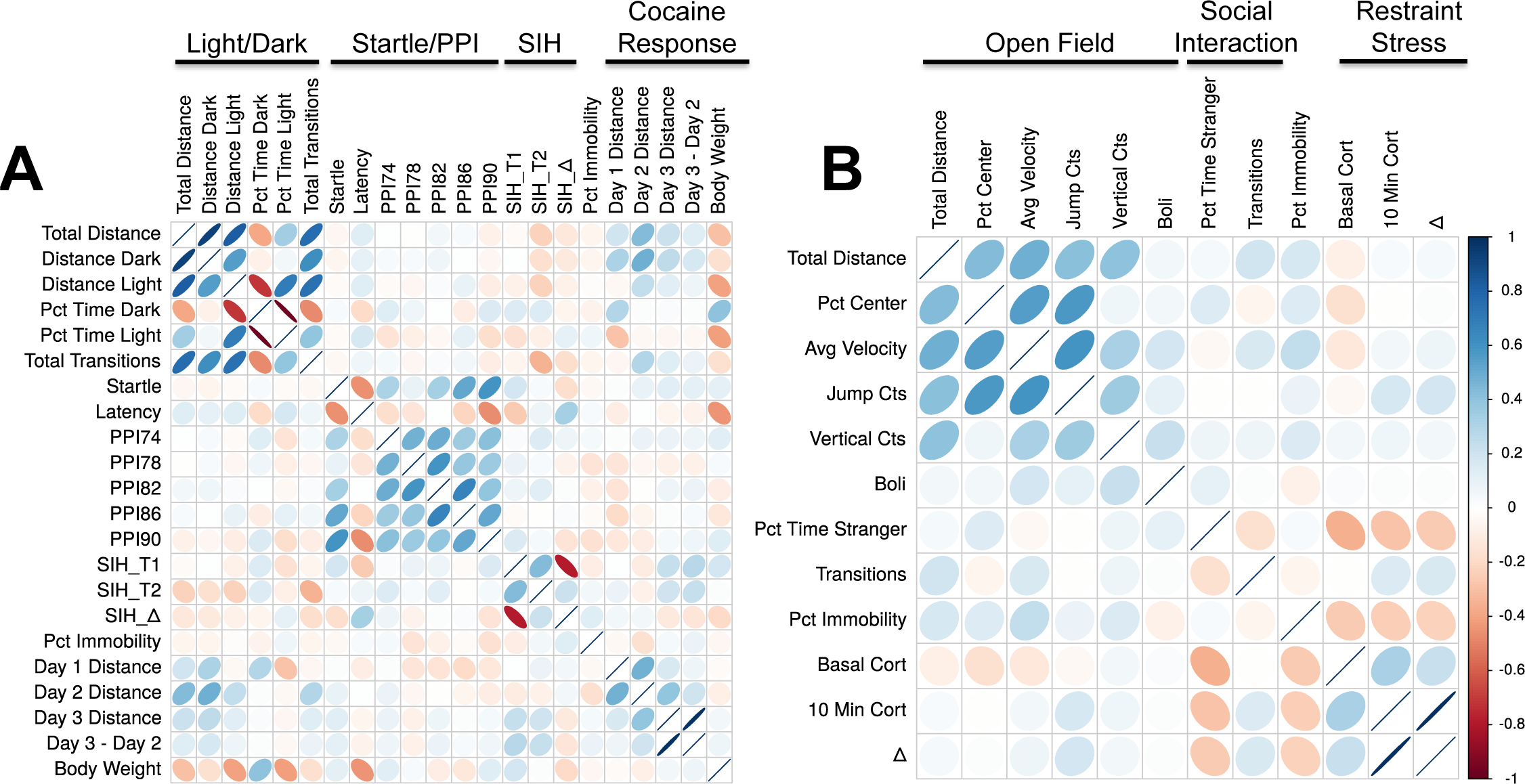
Correlation of behavioral phenotypes. A) Pipeline 1 behaviors. B) Pipeline 2 behaviors

**Figure S2.**
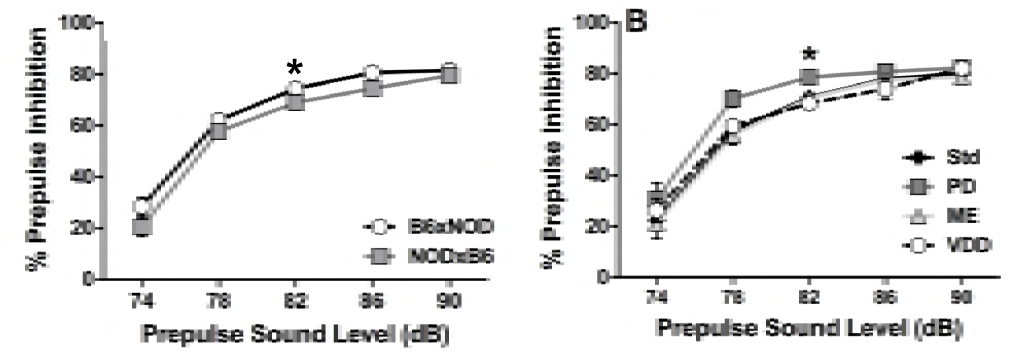
POE and perinatal diet effect on sensorimotor gating. (A) POE on sensorimotor gating; mean (bars indicate SEM) percent prepulse inhibition for B6×NOD (n=46) and NOD×B6 (n=45) mice. At 82 dB, a significant POE was observed on PPI (p=0.0307). (B) Diet effect on senorimotor gating; mean (bars indicate SEM) percent prepulse inhibition for standard (Std, n=31), vitamin D deficient (VDD, n=18), methyl enriched (ME, n=24) and protein deficient (PD, n=18) groups. At 82 dB, diet exposure significantly affected prepulse inhibition (p=0.00274).

**Figure S3.**
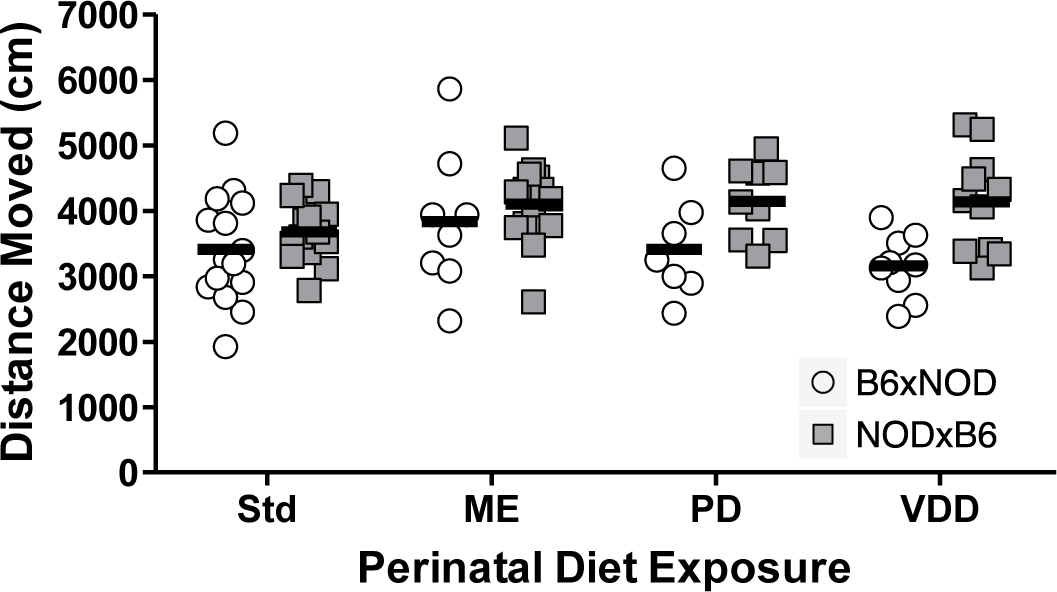
Non-significant but suggestive diet-by-POE on distance moved in a 10min open field test.) Data are individual B6×NOD and NOD×B6 animals exposed to Std (N=15,14), ME (N=8,14), PD (N=7,9) or VDD (N=9,11) diet. The pattern across diets follows that for percent center time (Figure 5).

**Figure S4.**
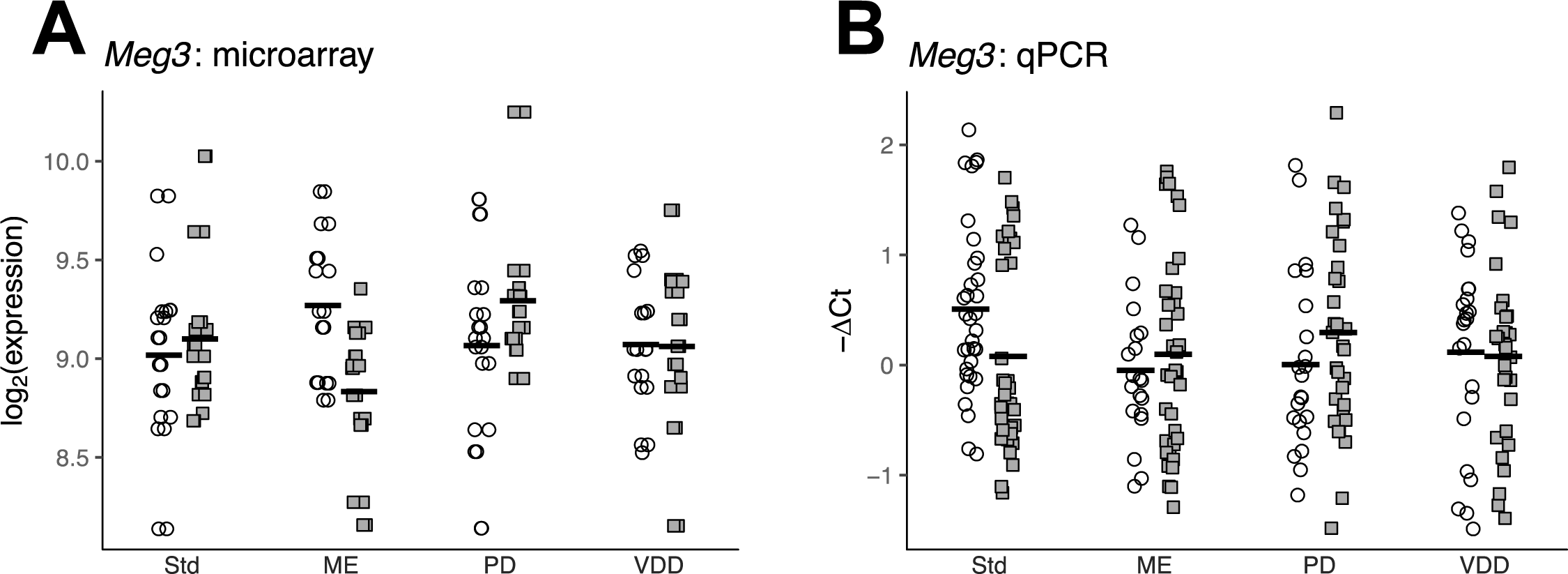
Perinatal diet-by-POE on Meg3 gene expression levels. Each point corresponds to *Meg3* expression of an individual B6xNOD or NODxB6 mouse exposed to standard (Std), vitamin d deficient (VDD), methyl enriched (ME), or protein deficient (PD) diet, as measured by A) microarray, and B) Taqman qPCR analysis. *Meg3* expression was significantly subject to diet-by-POE in the microarray analysis but not in qPCR validation.

**Figure S5.**
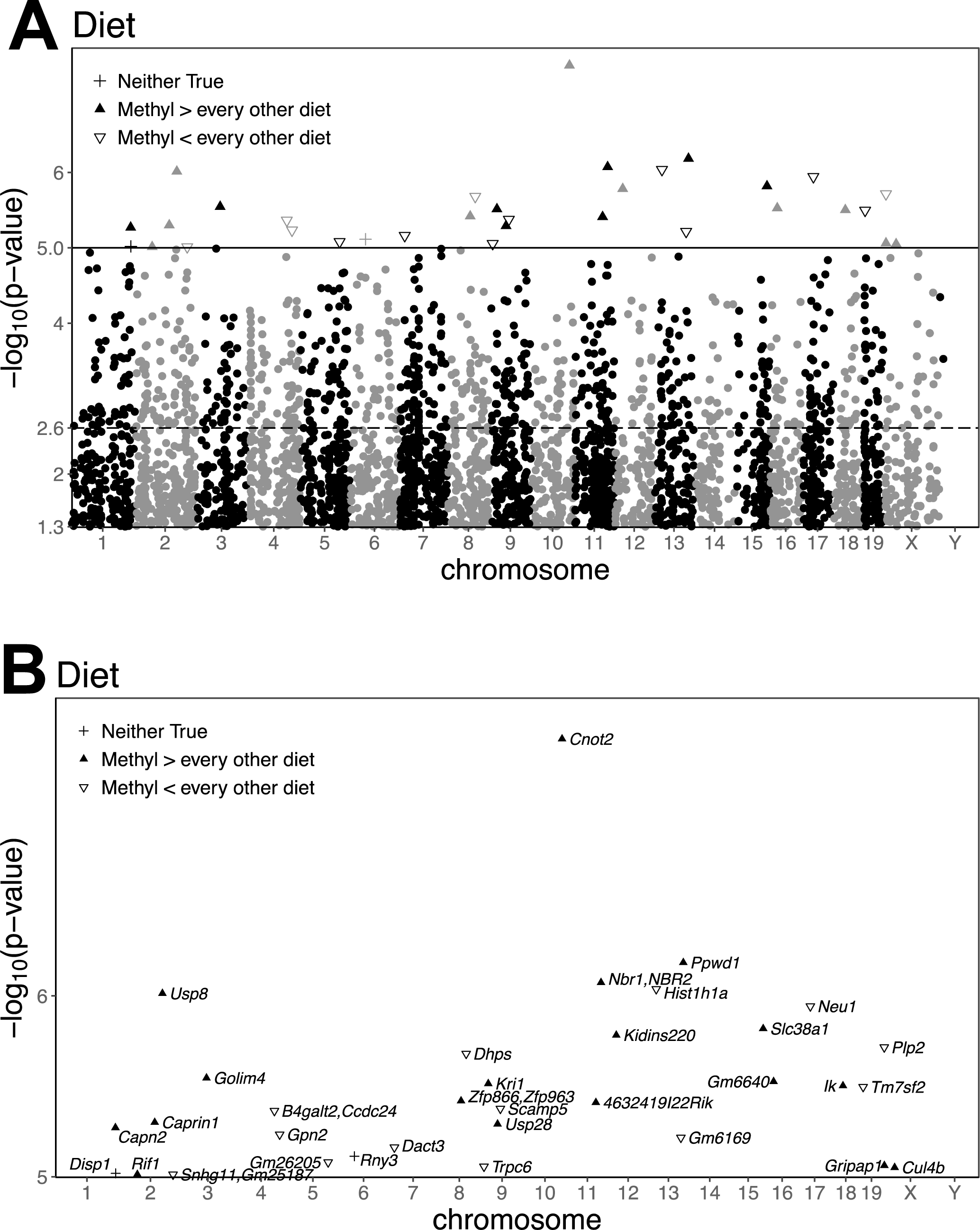
P-values of diet effects on gene expression. (A) Manhattan-like plot of p-values of diet effect on microarray-measured expression; each point corresponds to a probeset location and the p-value of the diet effect on that probeset’s expression. The dashed line represents the FDR threshold, and the solid line represents the FWER threshold. Probesets above the FWER threshold are marked by a shape, which depends on whether ME exposed mice have (on average) the highest expression relative to mice on the other diets (up arrow), the lowest expression relative to mice on the other diets (empty down arrow), or somewhere in middle of the 4 diets (plus sign); expression on the methyl diet is almost always at one or the other extreme. (B) Zoomed in view of just the genes significantly affected by diet. *Cnot2* is the most significantly affected

**Figure S6.**
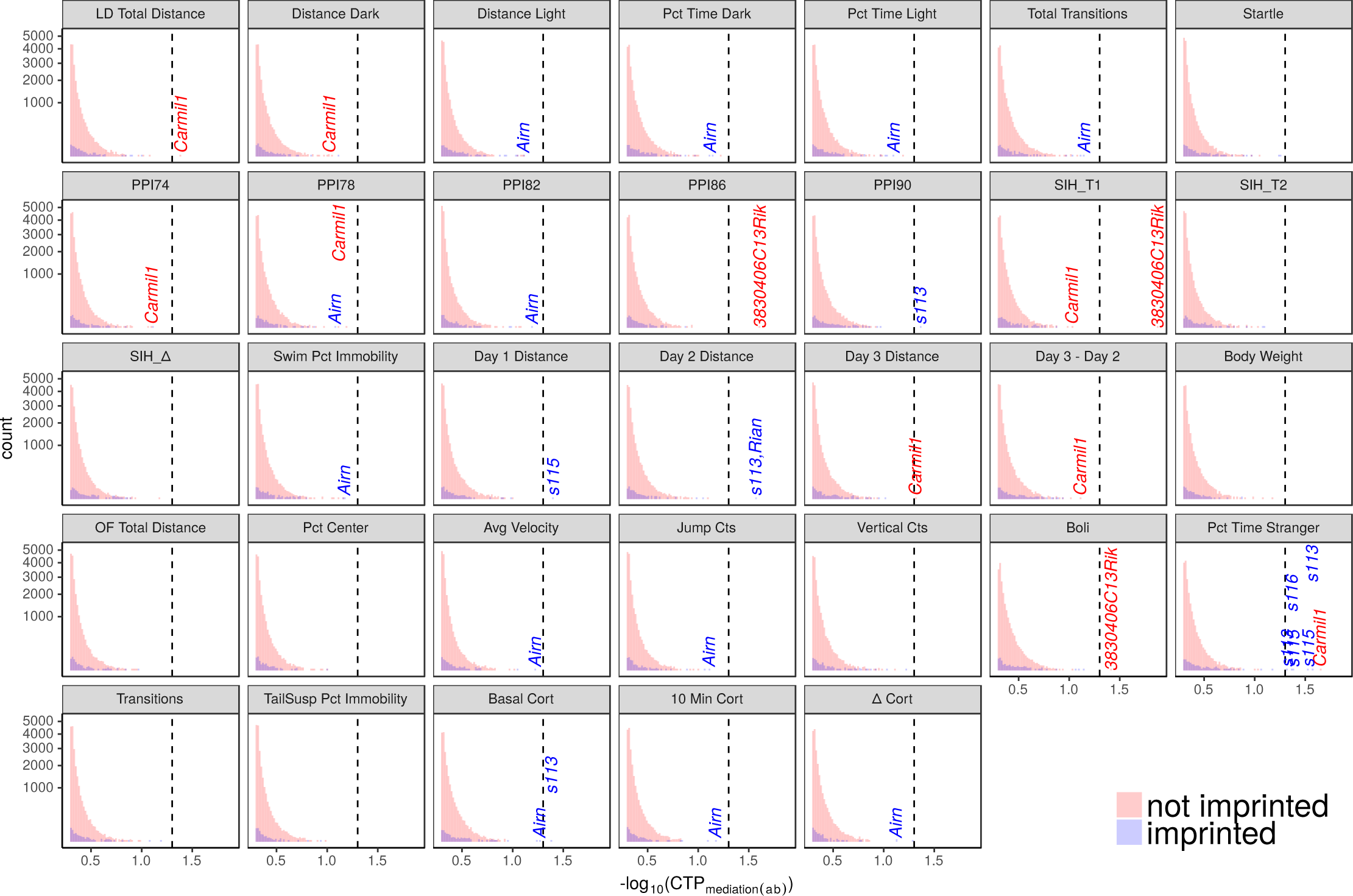
Histograms of the –log_10_ Combined Tail Probabilities (CTPs; akin to p-values) for candidate gene mediators of POE on various behaviors. Each panel corresponds to a behavioral outcome that may be mediated by gene expression. The red and blue histograms correspond to CTPs for non-imprinted and imprinted candidate mediators, respectively. Mediators having a CTP<.05 (threshold denoted by the dashed line) are labelled, as are *Airn* and *Carmil1* when they are one of the top-3 mediators. These two genes show up repeatedly (along with *Snord 113/115*) as one of the top mediators per behavior, especially when mediator/behavior pairs with a non-significant mediation p-value are also considered.

**Table S1.**
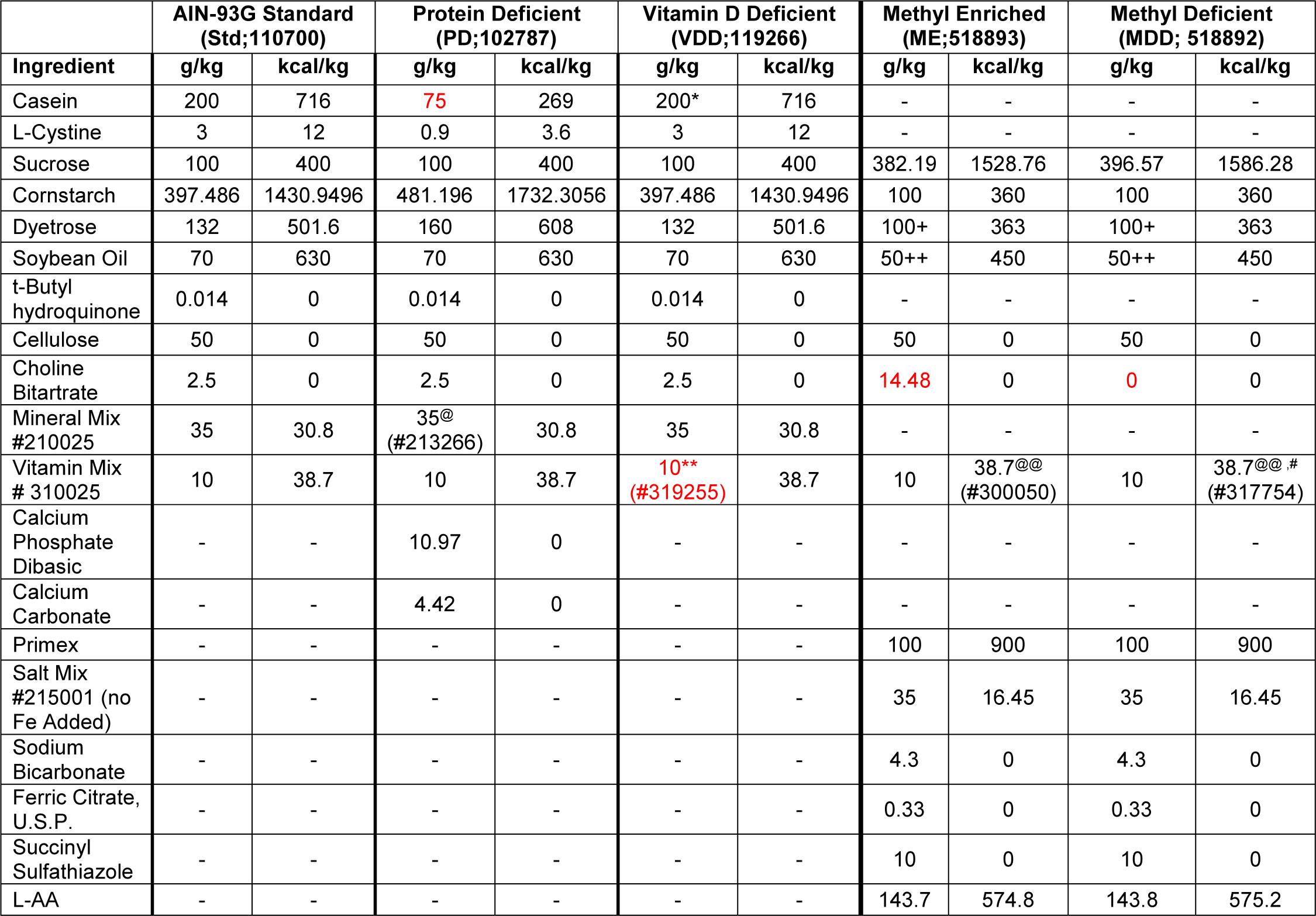
Nutritional content of experimental diets. The PD and VDD diets were nutritionally matched to the Std diet and the ME was matched to the MDD. Red indicates the main nutritional component that was changed in each diet (pelleted). The product number associated with each diet, vitamin mix, and mineral mix are provided (Dyets, Inc; Bethlehem, PA). @Ca and P free, @@Vitamin K1/Dextrose mix free w/ addition of menadione sodium bisulfite; #folic acid free; *vitamin free, ** no vitamin D, +dextrin instead of dyetrose, ++corn oil instead of soybean oil, #folic acid free

**Table S2.** Diets, number of dams, and female F1 hybrids per diet, broken down by various categories. The number of female F1 hybrids tested and the number of dams that produced those females in each behavior pipeline is broken down by reciprocal direction and perinatal diet exposure. The last column shows the breakdown of groups tested by behavior batch within each pipeline.

| Pipeline | Strain | Diet | # of Dams | # F1 Female Offspring | # Tested/Batch |  |  |
| --- | --- | --- | --- | --- | --- | --- | --- |
|  |  |  |  |  | 1 | 2 | 3 |
| 1 | B6xNOD | Standard | 8 | 20 | 5 | 6 | 9 |
|  |  | Methyl Enriched | 3 | 9 | 9 | 0 | 0 |
|  |  | Protein Deficient | 6 | 9 | 4 | 5 | 0 |
|  |  | Vitamin D Deficient | 3 | 8 | 0 | 4 | 4 |
|  | NODxB6 | Standard | 4 | 11 | 7 | 4 | 0 |
|  |  | Methyl Enriched | 4 | 15 | 6 | 9 | 0 |
|  |  | Protein Deficient | 3 | 9 | 2 | 3 | 4 |
|  |  | Vitamin D Deficient | 3 | 10 | 5 | 5 | 0 |
| 2 | B6xNOD | Standard | 5 | 15 | 5 | 6 | 4 |
|  |  | Methyl Enriched | 3 | 8 | 4 | 4 | 0 |
|  |  | Protein Deficient | 3 | 7 | 7 | 0 | 0 |
|  |  | Vitamin D Deficient | 3 | 9 | 0 | 3 | 6 |
|  | NODxB6 | Standard | 5 | 14 | 4 | 10 | 0 |
|  |  | Methyl Enriched | 3 | 14 | 9 | 5 | 0 |
|  |  | Protein Deficient | 4 | 9 | 4 | 5 | 0 |
|  |  | Vitamin D Deficient | 4 | 11 | 4 | 7 | 0 |

**Table S3.** Effect of perinatal diet and strain on breeding fitness. PND 21 = postnatal day 21 (time of weaning); F = females

| Dam Strain | Diet Exposure | # Dams Mated | # Litters | % Pregnant | # Pups Born | # Pups PND21 | % Survival PND21 | Mean Litter Size | # F Pups PND21 | % F Pups | # F Tested |
| --- | --- | --- | --- | --- | --- | --- | --- | --- | --- | --- | --- |
| C57BL/6J (B6) | Standard | 11 | 11 | 100 | 73 | 71 | 97.3 | 6.6 ± 2.4 | 35 | 49.30 | 35 |
|  | Methyl Enriched | 10 | 9 | 90 | 65 | 47 | 72.3 | 7.2 ± 1.9 | 21 | 44.68 | 17 |
|  | Methyl Donor Deficient | 12 | 2 | 16.7 | 18 | 0 | 0 | 9 | - | - | - |
|  | Protein Deficient | 12 | 12 | 100 | 52 | 44 | 84.6 | 4.3 ± 2.1 | 21 | 47.73 | 16 |
|  | Vitamin D Deficient | 10 | 10 | 100 | 54 | 52 | 96.3 | 6 ± 3 | 22 | 42.31 | 17 |
| NOD/ShiLtJ (NOD) | Standard | 7 | 7 | 100 | 63 | 61 | 96.8 | 9 ± 1.1 | 25 | 40.98 | 25 |
|  | Methyl Enriched | 8 | 7 | 87.5 | 63 | 52 | 82.5 | 9 ± 1.3 | 29 | 55.77 | 29 |
|  | Methyl Donor Deficient | 9 | 3 | 33.3 | 6 | 0 | 0 | 2 | - | - | - |
|  | Protein Deficient | 10 | 9 | 90 | 65 | 55 | 84.6 | 6.9 ± 1.7 | 20 | 36.36 | 18 |
|  | Vitamin D Deficient | 8 | 6 | 75 | 51 | 47 | 92.2 | 8.5 ± 2.1 | 21 | 44.68 | 21 |

**Table S4.**
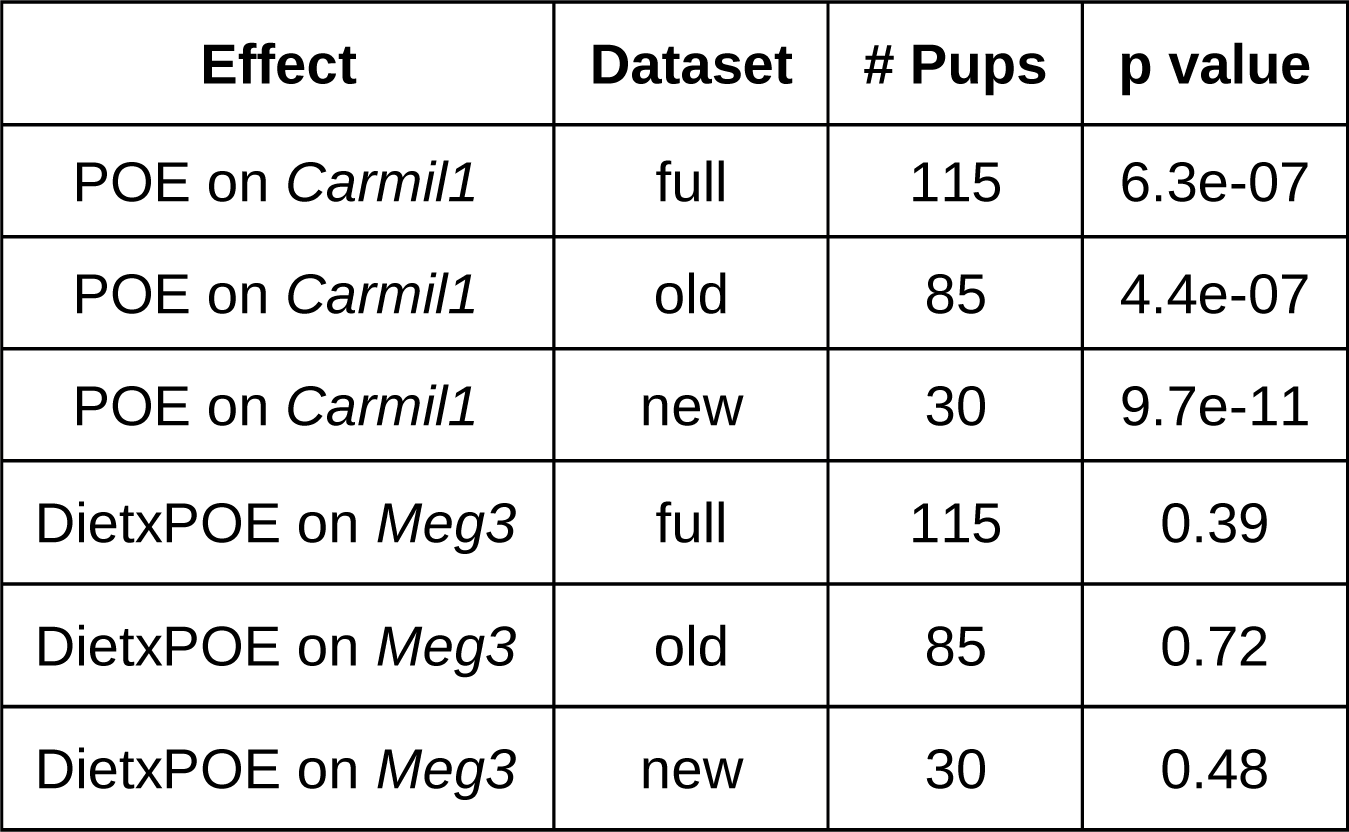
qPCR-based analysis of POE on *Carmil1* expression and Diet-by-POE on *Meg3* expression. Results shown for 3 datasets: i) mice that were both microassayed and qPCR’d (“old” data); ii) mice that were never microassayed but were qPCR’d (“new”); and iii) the union of the first 2 datasets (“full”). Parent-of-origin significantly affects qPCR-measured expression of *Carmil1* in all 3 datasets, whereas diet-by-parent-of-origin does not significantly affect qPCR-measured expression of *Meg3* in any dataset.

| Effect | Dataset | # Pups | p value |
| --- | --- | --- | --- |
| POE on *Carmil1* | full | 115 | 6.3e-07 |
| POE on *Carmil1* | old | 85 | 4.4e-07 |
| POE on *Carmil1* | new | 30 | 9.7e-11 |
| DietxPOE on *Meg3* | full | 115 | 0.39 |
| DietxPOE on *Meg3* | old | 85 | 0.72 |
| DietxPOE on *Meg3* | new | 30 | 0.48 |

**Table S5.**
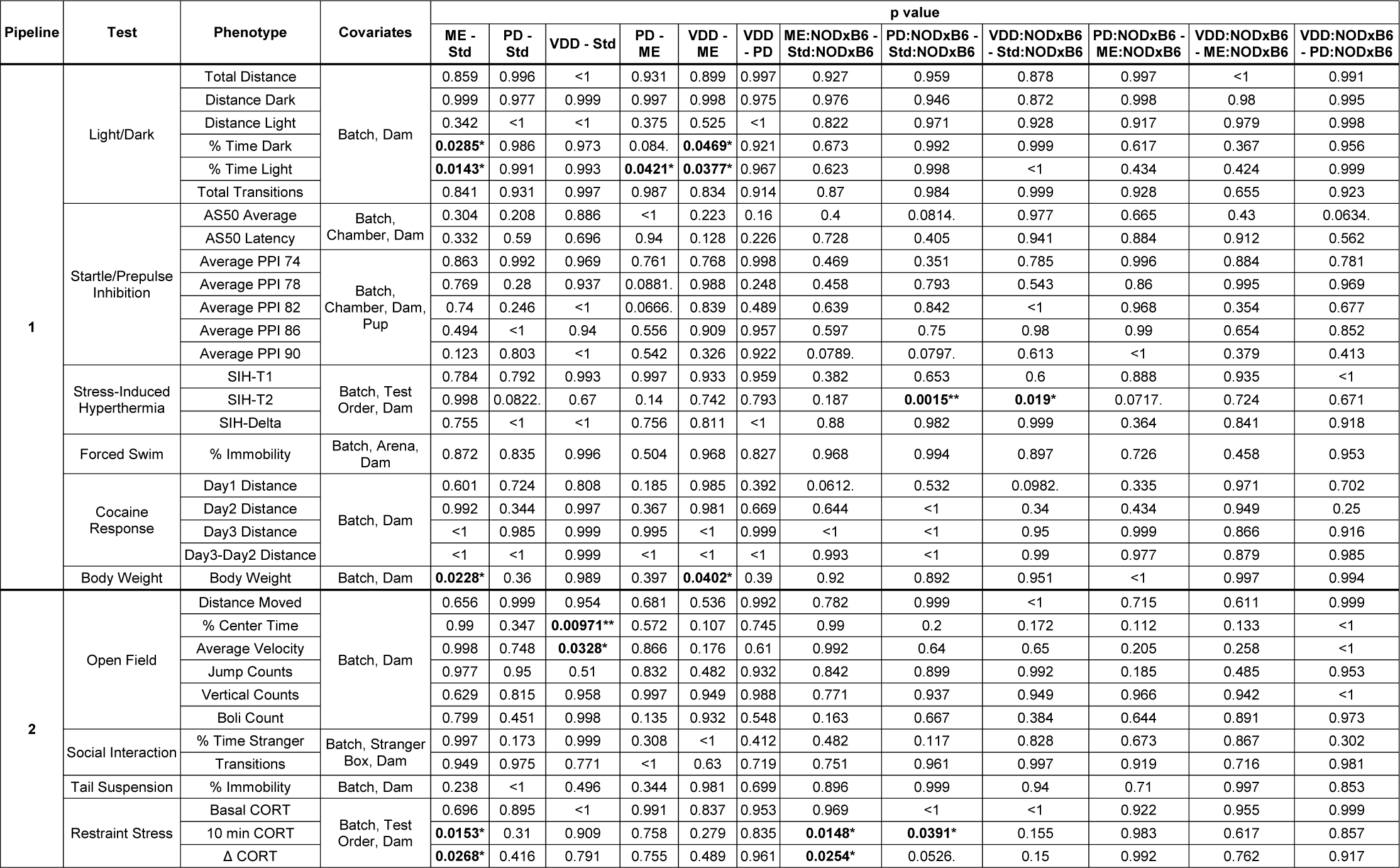
Tukey contrast p-values. For each phenotype, the table shows the modeled variables, along with the Tukey p-values for contrasts between all pairs of diets, and also between all pairs of diet-by-parent-of-origin effects. NOD:NODxB6 indicates an interaction between diet D with descent from maternal NOD. Significant values are **bolded**, and *, **, and ***, indicate significance levels of *0.05, **0.01, ***0.001 respectively. POE = parent of origin effect; PPI = prepulse inhibition; CORT = corticosterone; SIH-T1 = basal temperature; SIH-T2 = post-stress temperature; SIH-delta = (T2-T1)

**Table S6.** Coefficients and Combined Tail Probabilities (CTPs; akin to a p-value) for the significant gene mediators of *Carmil1* expression. The average mediation coefficient, a.k.a., the indirect effect, is *ab*, and is grouped together with its corresponding CTP. The direct effect, *c*’, is also grouped together with its associated CTP The mediation effect is generally *suppressing* the direct effect of POE; for 7 of 8 significant mediators, *ab* is opposite in sign to *c*’. *Airn* (as measured at probeset 10441787) is the most significant imprinted gene mediating the expression of *Carmil1.*

| Mediator Gene | Imprinted | Mediation Effect |  | Direct Effect |  | Suppressor |
| --- | --- | --- | --- | --- | --- | --- |
|  |  | ab | CTP | c' | CTP |  |
| 3830406C13Rik | FALSE | -0.465 | 0.00289 | 2.27 | <7.81e-05 | TRUE |
| *Irak1bp1* | FALSE | 0.158 | 0.00914 | 1.62 | <7.81e-05 | FALSE |
| *Airn_10441787* | TRUE | -0.166 | 0.0134 | 1.97 | <7.81e-05 | TRUE |
| *Tmem40* | FALSE | -0.115 | 0.0281 | 1.91 | <7.81e-05 | TRUE |
| *Pcdhb2* | FALSE | -0.123 | 0.0372 | 1.92 | <7.81e-05 | TRUE |
| *Mir485,Mirg* | TRUE | -0.116 | 0.0409 | 1.93 | <7.81e-05 | TRUE |

**Table S7.** Coefficients and Combined Tail Probabilities (CTPs; akin to a p-value) for the significant gene mediators of behavior. Mediation of POE was tested using each probeset’s expression as a mediator, against each behavior as an outcome; the behavior-probeset pairs in this table are the nominally significant associations. The mediator probeset is named according to the gene that is probed, followed by the specific probeset ID that was found to be significant if more than one probeset interrogates that gene. As was the case of mediation of expression, the mediation effect, a.k.a., the indirect effect, is *ab*, and is grouped together with its corresponding CTP. The direct effect, *c*’, is also grouped together with its associated CTP. As the coefficients are on a transformed scale, they are not especially informative, but they do demonstrate that for 17 of the 18 significant mediator-behavior pairings, the direct and indirect effect act opposite one another; i.e., when *ab* has the opposite sign of *c*’, mediation is suppressing the direct effect. We note that *Carmil1* and *Airn* both appear as significant mediators of behavior.

| Behavior | Mediator Gene | Imprinted | Mediation Effect |  | Direct Effect |  | Suppressor |
| --- | --- | --- | --- | --- | --- | --- | --- |
|  |  |  | ab | CTP | c' | CTP |  |
| LD Total Distance | *Carmil1* | FALSE | -1.39 | 0.0408 | 2.15 | 0.0393 | TRUE |
| Boli | 3830406C13Rik | FALSE | -0.857 | 0.0386 | 0.867 | 0.171 | TRUE |
| Day 1 Distance | *s115_10563911* | TRUE | 0.348 | 0.0398 | 0.304 | 0.339 | FALSE |
| Day 2 Distance | *s113,Rian* | TRUE | 0.66 | 0.0271 | -0.0562 | 0.478 | TRUE |
| Day 3 Distance | *Carmil1* | FALSE | -1.5 | 0.0486 | 1.62 | 0.0912 | TRUE |
| Basal Cort | *s113_10398354* | TRUE | 0.734 | 0.0415 | -1.24 | 0.12 | TRUE |
| SIH_T1 | 3830406C13Rik | FALSE | -1.43 | 0.0134 | 1.01 | 0.208 | TRUE |
| Pct Time Stranger | *Carmil1* | FALSE | 1 | 0.0227 | -0.658 | 0.297 | TRUE |
| Pct Time Stranger | *s113_10398354* | TRUE | 0.709 | 0.0272 | -0.526 | 0.317 | TRUE |
| Pct Time Stranger | *s115_10563949* | TRUE | 0.665 | 0.0296 | -0.432 | 0.366 | TRUE |
| Pct Time Stranger | *s115_10563959* | TRUE | 0.594 | 0.0415 | -0.381 | 0.377 | TRUE |
| Pct Time Stranger | *s116_10564209* | TRUE | -0.489 | 0.043 | 0.824 | 0.22 | TRUE |
| Pct Time Stranger | *s113_10398370* | TRUE | 0.567 | 0.0477 | -0.231 | 0.427 | TRUE |
| PPI86 | 3830406C13Rik | FALSE | 1.45 | 0.0245 | -1.86 | 0.0422 | TRUE |
| PPI90 | *s113_10398370* | TRUE | 0.633 | 0.0435 | -0.495 | 0.319 | TRUE |

**Table S8.** Genes that significantly mediate POE over all 34 behaviors in the aggregate; i.e., genes with a significant Combined Tail Probability<.05 (CTP), for POE mediation; the CTP essentially totals a gene’s Combined Tail Probabilities over every behavior. The table also includes the number of behaviors for which a given gene is among the 3 most significant mediators of POE (whether or not the POE on each behavior separately is significant), as well as the number of behaviors for which a given gene suppresses rather than contributes to POE. For example, *Carmil1* is one of the 3 most significant mediators of POE on 8 behaviors; for 26 behaviors, its mediation acts to suppress POE on that behavior; its CTP on POE mediation over all 34 behaviors is .00028. *S113/S115/S116* are shorthand for *Snord 113/115/116* respectively; Snord genes may be concatenated with a specific probeset ID when more than 1 probed region within the gene family is a mediator.

| Mediator Gene | Imprinted | # Behavior POEs |  | CTP |
| --- | --- | --- | --- | --- |
|  |  | strongly (top-3) mediated | suppressed |  |
| *Airn_10441787* | TRUE | 12 | 27 | 5.09e-05 |
| *Carmil1* | FALSE | 8 | 28 | 0.000518 |
| *s115_10563949* | TRUE | 7 | 23 | 0.000408 |
| *s115_10563911* | TRUE | 6 | 24 | 0.000857 |
| *1700063H04Rik* | FALSE | 4 | 18 | 0.0234 |
| *s113,Rian* | TRUE | 4 | 25 | 0.0152 |
| *3830406C13Rik* | FALSE | 4 | 24 | 0.00366 |
| *s113_10398370* | TRUE | 4 | 25 | 0.00484 |
| *s113_10398354* | TRUE | 4 | 27 | 0.0104 |
| *Mamdc2* | FALSE | 3 | 19 | 0.0208 |
| *Zrsr1* | TRUE | 3 | 19 | 0.0288 |
| *s116_10564209* | TRUE | 3 | 24 | 0.0192 |
| *Mir485,Mirg* | TRUE | 3 | 20 | 0.0164 |
| *Ndn* | TRUE | 2 | 19 | 0.00767 |
| *Irak1bp1* | FALSE | 1 | 18 | 0.0455 |
| *s115_10563915* | TRUE | 1 | 17 | 0.0214 |
| *Itga7* | FALSE | 1 | 24 | 0.0425 |
| *s115_10563989* | TRUE | 1 | 22 | 0.0324 |
| *Eps8l1* | FALSE | 1 | 22 | 0.0408 |
| *s115_10563959* | TRUE | 0 | 28 | 0.0259 |
| *3300002I08Rik* | FALSE | 0 | 24 | 0.0496 |

**Table S9** Genes whose expression is significantly affected by parent-of-origin, at the FWER threshold level. Chr = chromosome; N = no; Y = Yes; ME = methyl enriched diet; imprinted status determined using Crowley *et al.* (2015) and mousebook.org (Blake *et al.* 2010); p-values and q-values (FDR-corrected) are -log10 transformed.

**Table S10** Genes whose expression is significantly affected by perinatal diet exposure, at the FWER threshold level. Chr = chromosome; N = no; Y = Yes; ME Group rank = This gene’s expression rank, in mice exposed to methyl enriched (ME) diet, relative to mice on the other diets— *e.g.*, 1 means ME mice expressed this gene the most, whereas 4 means ME mice expressed this gene the least; imprinted status determined using Crowley *et al.* (2015) and mousebook.org (Blake *et al.* 2010); p-values and q-values (FDR-corrected) are -log10 transformed.

**Table S11** Genes whose expression is significantly affected by diet-by-POE, at the FWER threshold level. Chr = chromosome; N = no; Y = Yes; imprinted status determined using Crowley *et al.* (2015) and mousebook.org (Blake *et al.* 2010); p-values and q-values (FDR-corrected) are -log10 transformed.

